# Stimulus-induced theta band LFP oscillations format neuronal representations of social chemosignals in the mouse accessory olfactory bulb

**DOI:** 10.1101/2023.02.07.527460

**Authors:** Oksana Cohen, Anat Kahan, Idan Steinberg, Sebastian Malinowski, Dan Rokni, Marc Spehr, Yoram Ben-Shaul

**Author notes:** These authors contributed equally (Oksana Cohen, Anat Kahan).

## Abstract

Social communication is crucial for survival of many species. In most vertebrates, a dedicated chemosensory system, the vomeronasal system (VNS), evolved to process ethologically relevant chemosensory cues. The first central VNS stage is the accessory olfactory bulb (AOB), which sends information to downstream brain regions via AOB mitral cells (AOB-MCs). Recent studies provided important insights about the functional properties of AOB-MCs, but little is known about the principles that govern their coordinated activity. Here, we recorded local field potentials (LFPs) and single unit activity in the AOB while presenting natural stimuli to adult male and female mice. Our recordings reveal prominent LFP theta band oscillatory episodes with a characteristic spatial pattern across the AOB. We find that the AOB network shows varying degrees of similarity to this pattern throughout an experiment, as a function of sensory stimulation. Analysis of LFP signal polarity and single unit activity indicate that oscillatory episodes are generated locally within the AOB, likely representing a reciprocal interaction between AOB-MCs and granule cells (GCs). Notably, spike times of many AOB-MCs are constrained to the negative LFP oscillation phase, in a manner that can drastically affect integration by downstream processing stages. Based on these observations, we propose that LFP oscillations may gate, bind, and organize outgoing signals from individual AOB neurons to downstream processing stages. Our findings suggest that, as in other neuronal systems and brain regions, population level oscillations play a key role in organizing and enhancing transmission of socially relevant chemosensory information.

## Introduction

Social, reproductive, and defensive behaviors are crucial for survival, and in many organisms their implementation depends on chemical signaling and processing. In most vertebrates, including mice, this function is primarily served by the vomeronasal system (VNS), a compact but essential sensory system dedicated to processing chemical signals from other organisms (1–3). Although the VNS resembles the MOS, there are fundamental differences in their physiology and function. Studies in behaving (4) and anesthetized (5, 6) animals, and *ex vivo* preparations (7) examined spiking responses of neurons in the mouse AOB during sensory stimulation. These studies showed that AOB neurons display low baseline rates, and typically respond to chemical stimuli from conspecifics and non-conspecifics with rate elevations (8, 9). Anatomically, the first brain relay of the VNS, the AOB, appears like an integrative network, with its output neurons, AOB-MCs, sampling activity from multiple input channels (1, 10, 11). While individual vomeronasal ligands can elicit physiological and behavioral responses (12–16), most natural vomeronasal stimuli are complex bodily secretions, which contain numerous chemical components. Correspondingly, information about natural stimuli is likely represented by AOB-MCs in a distributed manner (5, 8, 9, 17, 18).

In the MOS, information about odorants is also conveyed to downstream regions by the concerted activity of ensembles of output neurons (19–23), and several processes were shown to organize and format output patterns of MOB neurons. These are often revealed by LFP signals, which typically reflect population-level activity (24–26). Specifically, a variety of olfactory bulb LFP oscillatory patterns, spanning the theta, beta, and gamma frequency ranges were described (27, 28). Although the exact mechanisms underlying these oscillations are only partly understood (29–34), it is known that they arise from both local (e.g., within the MOB) and global interactions (e.g., involving multiple olfactory brain regions), and are directly influenced by breathing. Together, these oscillatory patterns reflect processes that may gate, bind, and convey parallel streams of information (i.e., multiplex), as it is transmitted from the MOB to downstream regions. More specifically, activity of neurons with respect to each other, or with respect to the breathing cycle, can be informative about the stimulus (22, 23, 35–42) suggesting that they are important for chemosensory information processing. Yet, similar phenomena were not reported in the VNS.

One of the most notable differences between the MOS and the VNS involves stimulus uptake. In the MOS, stimulus uptake is linked to breathing, and active sampling modifies sniffing behavior and hence the manner by which information is relayed to downstream regions (43–45). In contrast, VNS stimulus uptake is largely uncoupled from breathing, and instead relies on active suction by the vomeronasal organ (5, 46, 47). Other differences between the systems include the intrinsic properties of olfactory bulb neurons (48–50), their interactions with each other (2, 10, 51), and the properties of downstream regions that receive the information (23, 35, 42, 52). Together, these observations imply that if, like the MOS, VNS information processing involves coordinated patterns of population activity, they are likely to differ in their phenomenological expression and underlying mechanisms.

Previously, we and others (53–56) have identified infra-slow oscillations in the AOB. In addition, a number of studies have shown a broad range of LFP AOB oscillations that can reflect experience, context, and potentially active arousal/sampling (57–61). However, explicit relationships between these rhythmic population activity patterns and sensory stimulation and, more importantly, their potential interaction with single unit spiking activity are unknown.

Here, we describe prominent theta band LFP oscillations in the AOB. We show that they are induced by stimulus presentation and that their magnitude is stimulus-dependent. Notably, in addition to full-blown oscillations, the AOB network also displays a more subtle pattern of stimulus-dependent activity that closely resembles the oscillations in its spatial distribution. Our analysis suggests that once sensory stimulation is sufficiently intense, these latent patterns develop into prominent oscillations. Based on their spatial and temporal features, we propose that these LFP patterns reflect a periodic exchange within the AOB network, likely involving AOB-MCs and GCs. Most importantly, we show that these oscillations constrain spike times of individual AOB neurons and can therefore serve as a thresholding, amplifying, and potentially also multiplexing mechanism. Thus, despite prominent differences in their constituent elements and connectivity, it appears that, like the MOS, the VNS includes circuit elements that produce population level oscillations, which serve to orchestrate information transfer to downstream regions.

## Results

To investigate AOB LFPs during sensory processing, we presented vomeronasal stimuli to anesthetized mice by direct application to the nostril, and electrically activated the VNO (**Fig. 1A**). In a typical experiment, each stimulus is presented repeatedly (typically 4-5) in a pseudorandom order (62). During stimulus presentation, we record both LFP (0.5 – 300 Hz) and single unit activity using planar multielectrode arrays, which sample a substantial extent of the AOB. Our dataset contains 34 recordings sessions from adult male and female mice. Details about each session are given in **Table S1**. We first describe recordings made with 32-channel probes (**Table S1**), advanced in parallel to the AOB external cellular layer (ECL) (**Fig. 1A-C**), maximizing the number of contact sites within the ECL, which contains AOB-MC cell bodies (10).

**Fig. 1.**
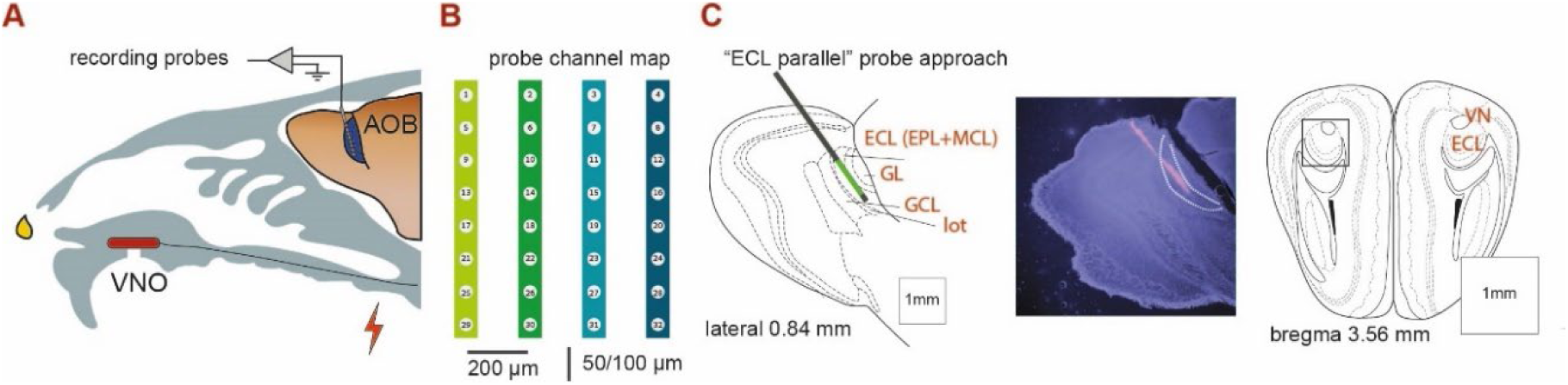
Experimental setup and recording sites. **A.** Experimental setup showing electrode array. The lightning bolt indicates stimulation of the sympathetic nerve trunk which innervates the VNO. During each trial, a drop of stimulus (2 μl) is applied to the nostril (application event) and, after 20 s, activate the VNO via a cuff electrode (stimulation event) placed around the sympathetic nerve trunk. After 40 s, the nasal cavity is flushed and another stimulus is presented. See also Fig. 2A. **B.** Schematic of the 4×8 recording array. Inter-site distances along a shank are 50 or 100 μm (see **Table S1**). **C.** Atlas images of the AOB in sagittal (left) and coronal (right) views. Images are adapted from (63). The black line in the sagittal section shows the approximate orientation of the recording probe. The green line shows the approximate extent of the recording region along one shank (with 100 μm inter-site spacing). Histological image shows the path of a DiI-stained probe (red), with nuclear DAPI counterstaining (blue). The broken white lines delineate the AOB ECL. The square within the coronal section shows the approximate horizontal extent of the probes (600 μm). ECL: external cellular later. EPL: external plexiform layer. MCL: mitral cell layer, GL: Glomerular layer. GCL: Granule cell layer. Lot: lateral olfactory tract, VN: vomeronasal nerve layer.

### The AOB displays a spatially stereotypical pattern of oscillations

To detect prominent LFP patterns, we first used a custom written data browser and manually scanned the multichannel data. Occasionally, we observed distinct oscillatory events that appeared across a subset of channels at varying amplitudes (**Fig. 2A, B**). As detailed below, these oscillatory episodes vary in frequency (both within and across an episode), but broadly fall within the theta range (2-12 Hz). Notably, within a session, all manually detected oscillatory events shared a similar across-channel envelope pattern (**Fig 2C, D, Fig. S1**). When LFP signals from each channel are plotted according to their relative spatial locations, it is apparent that regions with high LFP activity correspond to a central active area, surrounded by a region with considerably lower magnitudes. This is shown for one oscillatory event in **Fig. 2E**, and for the average envelope power across all manually detected events from the same session in **Fig 2F**. As electrode positioning within the AOB is variable, these envelope distributions are distinct for each recording session, yet, almost all reveal a central hotspot of activity. Examples from different sessions are shown in **Figs. 2G** and **Fig. S2**. Considering AOB anatomy and probe dimensions (**Fig. 1C**), these patterns imply that signals are generated locally, and do not represent volume conduction from other regions. Further evidence that supports local signal generation is provided in later sections.

**Fig. 2.**
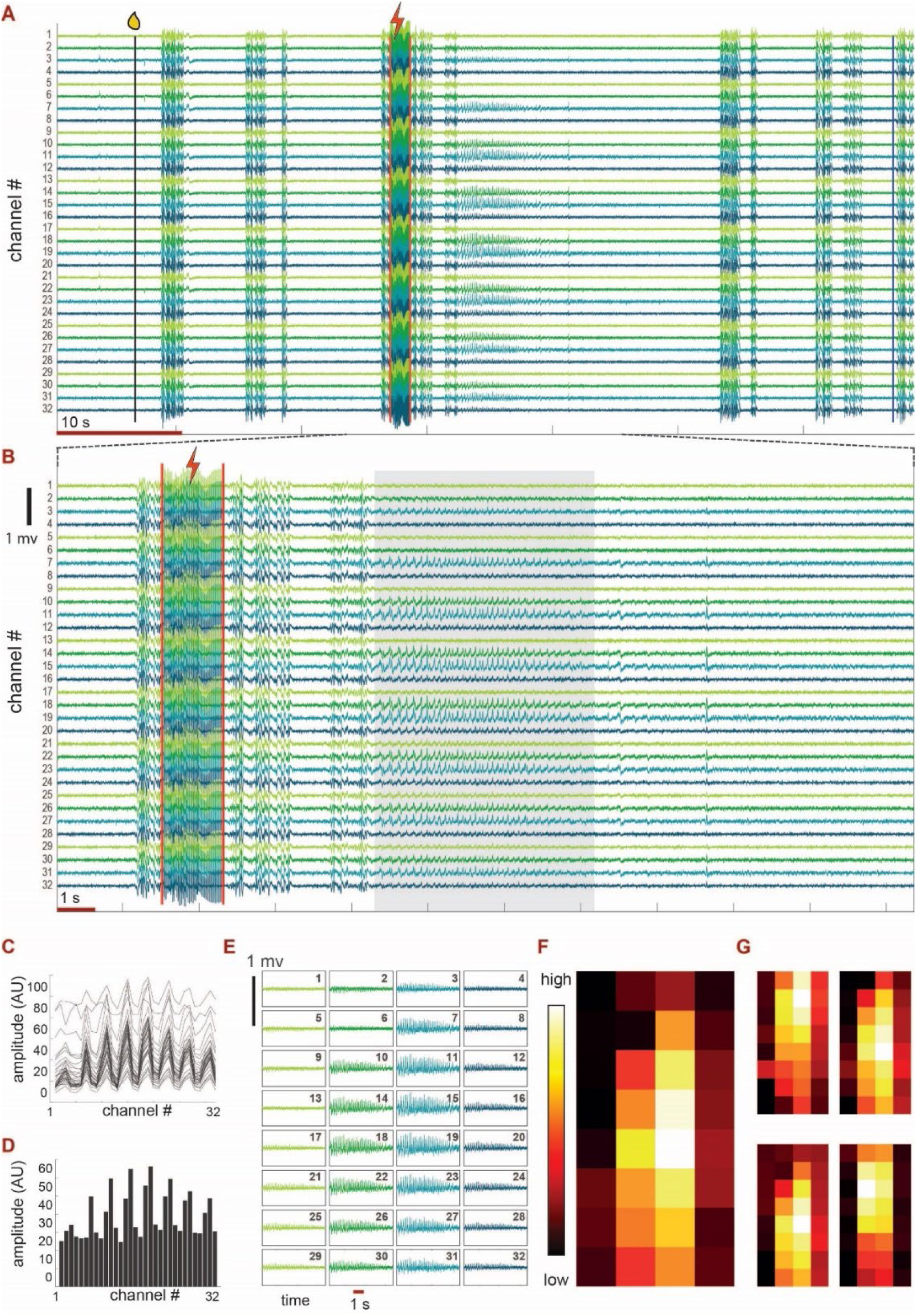
Basic analysis of the LFP data. **A.** LFP signals from 32 channels during one trial (∼70 s). Stimulus *application* is indicated by the black vertical line and drop icon, VNO *stimulation* is indicated by the red vertical lines and the lightning bolt icon, and the rightmost vertical blue line represents the beginning of the wash period between trials. Channels are arranged according to their serial numbers (not spatial positions). Channel colors indicate the shank of the recording site (see Fig. 1B). LFP amplitudes are normalized to fit within the plot. A vertical distance of 1 (i.e., channel number baselines) corresponds to 357 µV. **B**. Magnified view of a section shown in panel A. **C.** LFP envelopes (across all channels) associated with all manually defined events for the same session from which A and B are taken. **E.** The mean LFP envelope across all events in D. **E.** A section containing the oscillatory episode in B, with channels plotted according to their physical arrangement. In this session, horizontal distances (between shanks, columns) are 200 µm and vertical distances among sites are 100 μm. Channel numbers are indicated in each panel and correspond to the numbers in panels A-B and Fig. 1B. **F.** Mean LFP envelope (across all manually defined oscillatory events) drawn spatially as a heat map (scaled to show the entire range). The leftmost image corresponds to the template in E. **G.** Mean LFP envelopes for four other sessions (see also **Fig. S2**).

### Oscillatory episodes comprise frequency sweeps within the theta range

We began by analyzing the spectro-temporal characteristics of the oscillatory episodes. The procedure for identification, temporal, spectral, and phase characterization of episodes is described in Methods (see **Fig. S3**). At the end of this procedure, we have, for each oscillatory event, temporally-evolving frequency and phase estimates (used in the next section). These estimates are based on the first time-frequency “ridge” of the synchrosqueezed wavelet transform (see Methods for rationale and further details). We make a distinction between “complete” episodes (n = 89), which contain an entire oscillation event from start to end without interruption, and all other episodes (n = 2254) which contain a dominant theta frequency component, but do not contain the entire event. **Fig. 3A** shows examples of twelve complete episodes from two recording sessions. These examples underscore the diversity in episode duration and frequency content. Similar plots for all complete episodes are shown in **Fig. S4**. Examination of all episodes shows that while many episodes follow a smooth up and down sweep in the frequency domain, others assume a more complex structure. Although individual events vary in their mean frequencies, defined as the amplitude weighted mean frequency over the entire episode (see Methods), all fall within the broad theta range (**Fig. 3B**, mean of mean frequencies: 5.47 Hz). As shown in these examples and in **Fig. S4**, episodes also vary in duration (**Fig. 3C**). Although most episodes are shorter than 5 s, some are over 15 s long. The distribution of episode durations and frequencies for all episodes (n = 2343. mean, mean frequency 7.1 Hz, mean duration 2.29 s) are shown in **Fig. S5E**. In conclusion, although oscillatory episodes fall within the theta range, they vary in both duration and spectral characteristics. Notably, there are also differences between episodes recorded during a single session, indicating that variability is not merely due to different experimental preparations or electrode positions.

**Fig. 3.**
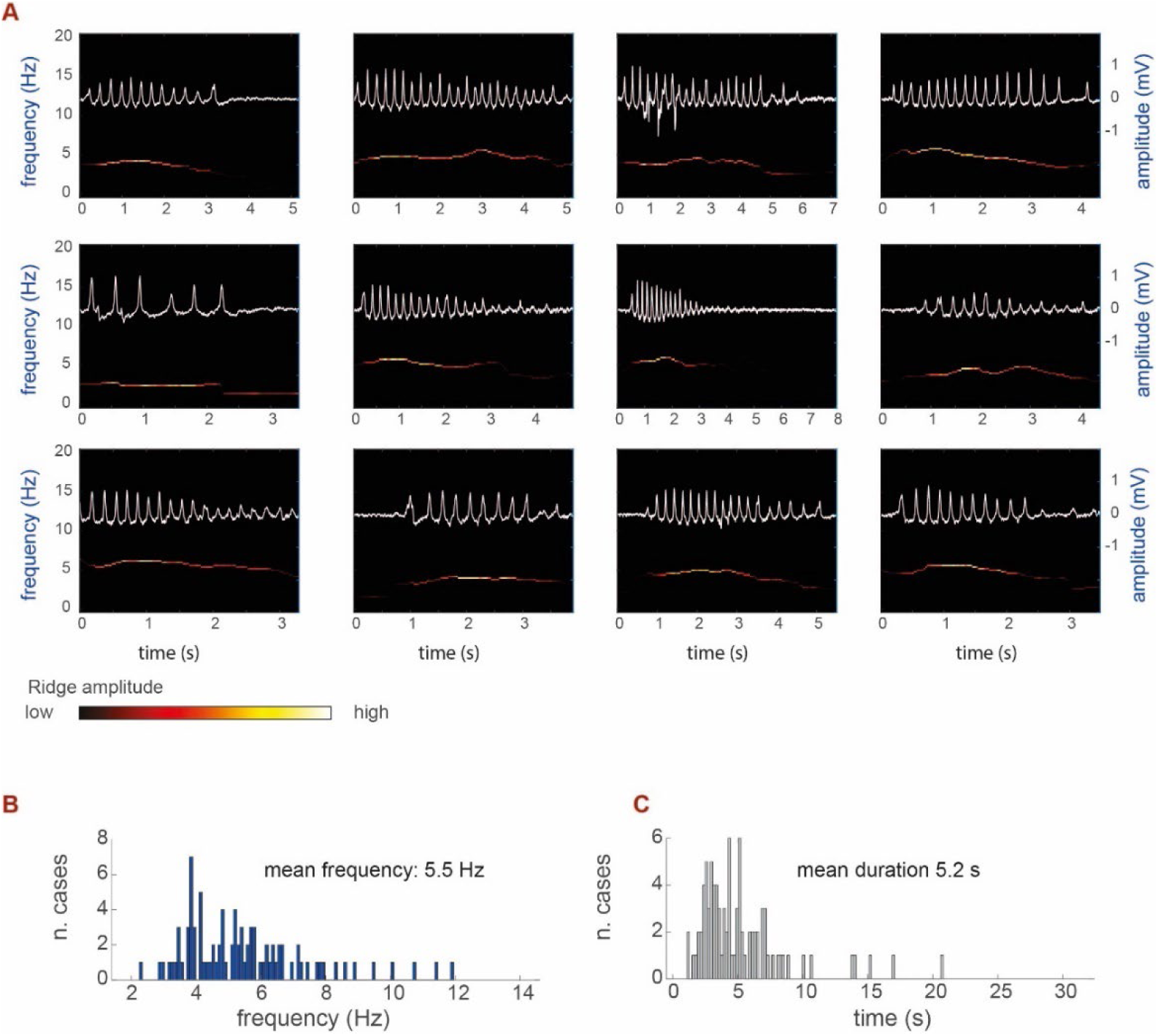
Oscillatory episodes vary in duration and spectral structure. **A.** Spectral analysis of twelve oscillatory episodes. The white traces show the original signals, with amplitude scale indicated on the right side. The colored trace is a spectrogram showing the time evolving frequency (see Methods). The same vertical axes are used for all episodes, but horizontal (time) scales vary. Note that many of the episodes shown an increase and then a decrease in frequency. All episodes on the top row and the left episode on the bottom row are from the same session. All the other episodes are from another session. **B-C.** Distribution of mean frequencies (B) and durations (C) for all complete episodes (n = 89).

### Spiking activity is locked to the LFP phase

In various brain regions and systems, and the MOS in particular (27), LFP oscillations can reflect orchestrated action potential timing of individual neurons (26). For example, in the MOB, constrained spike timing can facilitate activation of downstream brain regions, or multiplex information in individual channels (23, 34, 39, 40, 64). To investigate whether a similar phenomenon occurs in the AOB, we studied the relationship between spike timing and the LFP phase. As mentioned above (Methods), the frequency and phase estimates are based on the first time-frequency ridge of the synchrosqueezed wavelet transform. To determine the instantaneous LFP phase at each time point, we applied the Hilbert transform to the first ridge (see Methods and **Fig. S3**). We only consider oscillatory episodes for which the first ridge provided an above threshold reconstruction of the original signal (reconstruction score > 0.5, see Methods), and for which the mean frequency is within the theta range (2-14 Hz). importantly, the phase is derived from the channel with the highest LFP energy. This could be a concern if different channels displayed varying phase lags with respect to each other, so that a single channel would not be representative of all. However, our analysis indicates that during oscillatory episodes, the phase is aligned across all channels (**Fig. S6**, see also **Fig. 7**).

Having determined the phase of each action potential, we can address several hypotheses. Below, we compare the distributions obtained with the real data to those obtained with phase-shuffled data (see Methods). First, we tested whether AOB spiking activity depends on the LFP phase. To this end, we tested if the population of single units displays phase distributions that deviate from uniformity, using the omnibus test for circular data (65). **Fig. 4A** shows examples of phase distributions (green bars) of eight single units with significant deviations from uniformity. Under the null hypothesis that spike timing is independent of the LFP phase, the distribution of p-values (for deviations from uniformity) across the population should itself be uniform, and the cumulative distribution should follow the diagonal. As shown in **Fig. 4B**, this is not the case. There is a marked excess of small p-values for the real, but not for the shuffled data. The probability of obtaining such an excess of small p-values (e.g., smaller than 0.05) can be derived from the binomial distribution (see Methods), and is very small (n = 331 single units, probability: 3.9 •10^-36^). The corresponding p-value distribution for the phase-shuffled data is uniform, without evidence for excess of small p-value (binomial p = 0.90). We note that (as expected) shuffled p-values slightly change each time the analysis is run, but our conclusions remain entirely valid across many repeated runs. Thus, we first conclude that, as a population, AOB-MCs display spike time preference with respect to the LFP phase.

**Fig. 4.**
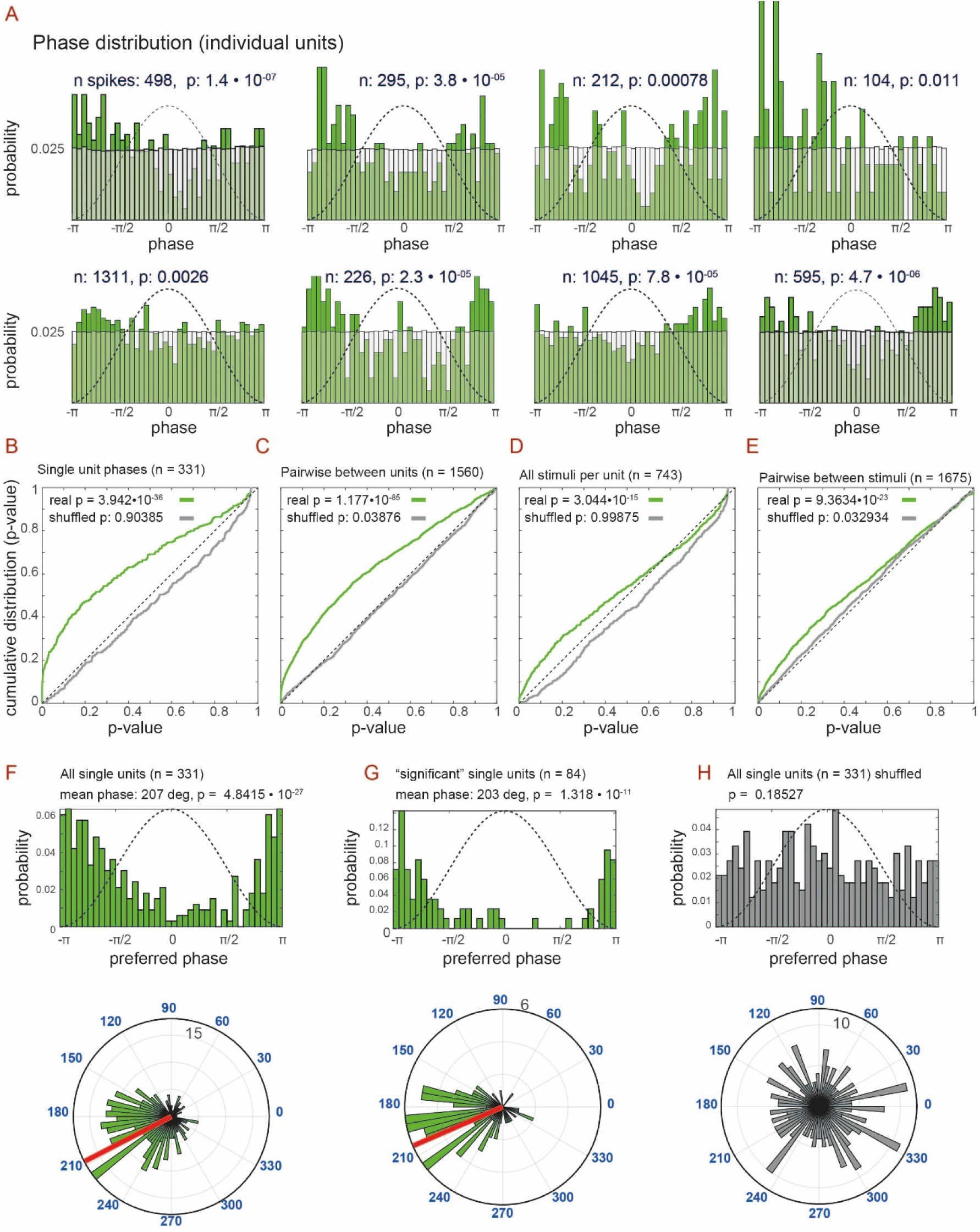
The distributions of single unit spike times relative to the oscillation phase are nonuniform. **A.** Phase histograms of eight single units; number of spikes and p-value for tests of uniformity are indicated in each panel. Broken line represents the LFP cycle for reference. Green bars show the units’ phase distribution. Lighter bars show the distributions of the LFP phases across all episodes during the corresponding recording session, and are flat as expected. **B.** Cumulative distribution of p-values for the test of uniformity (of spike times relative to LFP phase) across all single-units. Green and gray traces correspond to real and shuffled phase data, respectively. **C.** Cumulative distribution of p-values for pairwise comparison of phase medians across all simultaneously recorded unit pairs. **D**. Cumulative distribution of p-values for each unit and stimulus combination. **E.** Cumulative distribution of p-values for pairwise comparison of phase medians across all stimulus pairs for each unit. For the analysis in **D-E** we only consider unit-stimulus combinations with at least 50 spikes. **F.** Preferred phase distributions across all single-units and **G**. single units with a significant deviation from phase uniformity (p < 0.05), in both linear (top), and polar (bottom) histograms. The red bars in the polar plots show the mean phase. Broken lines in linear plots shows the LFP cycle for comparison. **H:** Like panel F, but for phase shuffled data.

Next, we consider the distribution of the units’ preferred phases (e.g., circular means). Under one extreme scenario, preferred phases homogeneously tile the LFP cycle period, resulting in a uniform distribution of preferred phases. At the other extreme, all units share similar preferred phases. The actual distribution (**Fig. 4F**) again shows a marked deviation from uniformity (N = 331, mean phase: 207 degrees, omnibus test for uniformity p = 4.8•10^-27^). Very similar results are obtained when only single-units with a significant deviation from uniformity are considered (**Fig. 4G**, n = 84, probability: 1.3•10^-11^ mean phase: 203 degrees). Notably, preferred phases correspond to the negative LFP phase, consistent with a current sink in the ECL. With shuffled data (**Fig. 4H**), preferred direction distributions do not deviate from uniformity.

Importantly, pairwise comparisons of phase distributions between simultaneously recorded units, using a non-parametric multi-sample test for equal medians (65), shows that many unit-pairs show distinct phase distributions (**Fig. 4C**). Specifically, the analysis reveals a marked excess of significant differences (n = 1560 unit-pairs, probability to obtain such an excess of p-values < 0.05 under the binomial model: 1.18•10^-85^, for the shuffled data: p = 0.038). Although the probability for the shuffled data is below 0.05, the corresponding probability for the real data is orders of magnitude smaller (see Methods for further discussion). We therefore conclude that along with the general preference for the negative phase of the LFP cycle, there are also frequent differences between the specific phase preferences of individual units.

Finally, we test a more elaborate hypothesis, according to which individual units show distinct phase distributions when different stimuli are presented. First, we consider the population of unit-event pairs and ask, as before, whether there is an excess of significant p-values (under the null hypothesis of uniform distributions). This is true for the real data (n = 743 unit-event pairs, probability under the binomial model: 3.04•10^-15^, **Fig. 4D**), but not for shuffled data (p = 0.99). Next, we compare phase distributions of the same unit during presentation of different stimulus pairs. As shown in **Fig. 4E**, there is a marked excess of significant pairwise differences for the real data (1675 comparisons, probability under the binomial mode: 9.36•10^-23^), but not for the phase-shuffled data (p = 0.03). This analysis is thus consistent with the idea that a given unit can follow distinct phase distributions, depending on the sensory stimulus.

Taken together, we have shown that individual neurons in the AOB have a preferred firing phase within the LFP theta cycle, and that the preferred phase varies among neurons, and is stimulus dependent. This temporal structure of firing can play important roles in information transfer to downstream regions. We further address the potential implications of these findings below and the Discussion.

### Oscillatory periods represent an extreme expression of a continuously varying network state which reflects sensory stimulation

Recall that all oscillatory episodes within a session are characterized by a stereotypic energy distribution across recording sites. We denote this as the *oscillating network energy distribution* (*ONED,* see **Fig 2F, G** and **Fig. S2**). The ONED represents a particular network state, and next we study its expression throughout the experiment. Specifically, we ask whether the network resembles the ONED only during oscillatory periods, or rather, that it represents a graded pattern throughout the experiment. Furthermore, we asked whether the ONED is corelated with stimulus presentation, as might be the case if it plays a role in information processing. We use the term *template match signature* (TMS), to denote this measure of similarity to the oscillating state (ONED). To derive the TMS, we calculate the linear correlation coefficient between the ONED template and the LFP envelope across all channels at each point in time. The result is smoothed, rectified, and sharpened to increase contrast, yielding a measure of similarity between ongoing network activity and the oscillatory template (see **Fig. S3A** and Methods).

If the ONED is related to vomeronasal sensory processing, the TMS should be modulated by sensory stimulation. **Fig. 5A** shows an example of the TMS during an entire recording session, with epochs of stimulus presentation denoted by green horizontal bars. As illustrated in this representative example, the TMS increases during virtually every period of stimulus presentation. To analyze the mean effect of stimulus presentation on the TMS, we derived its stimulus-triggered average (staTMS). For these analyses, we focus on one stimulus set, which includes three stimuli, each at three different dilutions (set D2 in **Table S1**). The global staTMS, based on all trials from all sessions is shown in **Fig. 5B**. The staTMS reveals that the TMS is tightly linked to stimulus presentation, first increasing during stimulus *application*, and further rising following electrical VNO *stimulation*. We note that this temporal profile is fully consistent with the observations that AOB neurons sometimes respond after stimulus *application*, prior to electrical activation of the sympathetic nerve trunk (66, 67). Very similar results are obtained in another set of experiments using different stimuli (**Fig. S7A**). In summary, these analyses indicate that the TMS, and hence the oscillating network state are closely correlated with chemosensory stimulus processing in the AOB.

**Fig 5.**
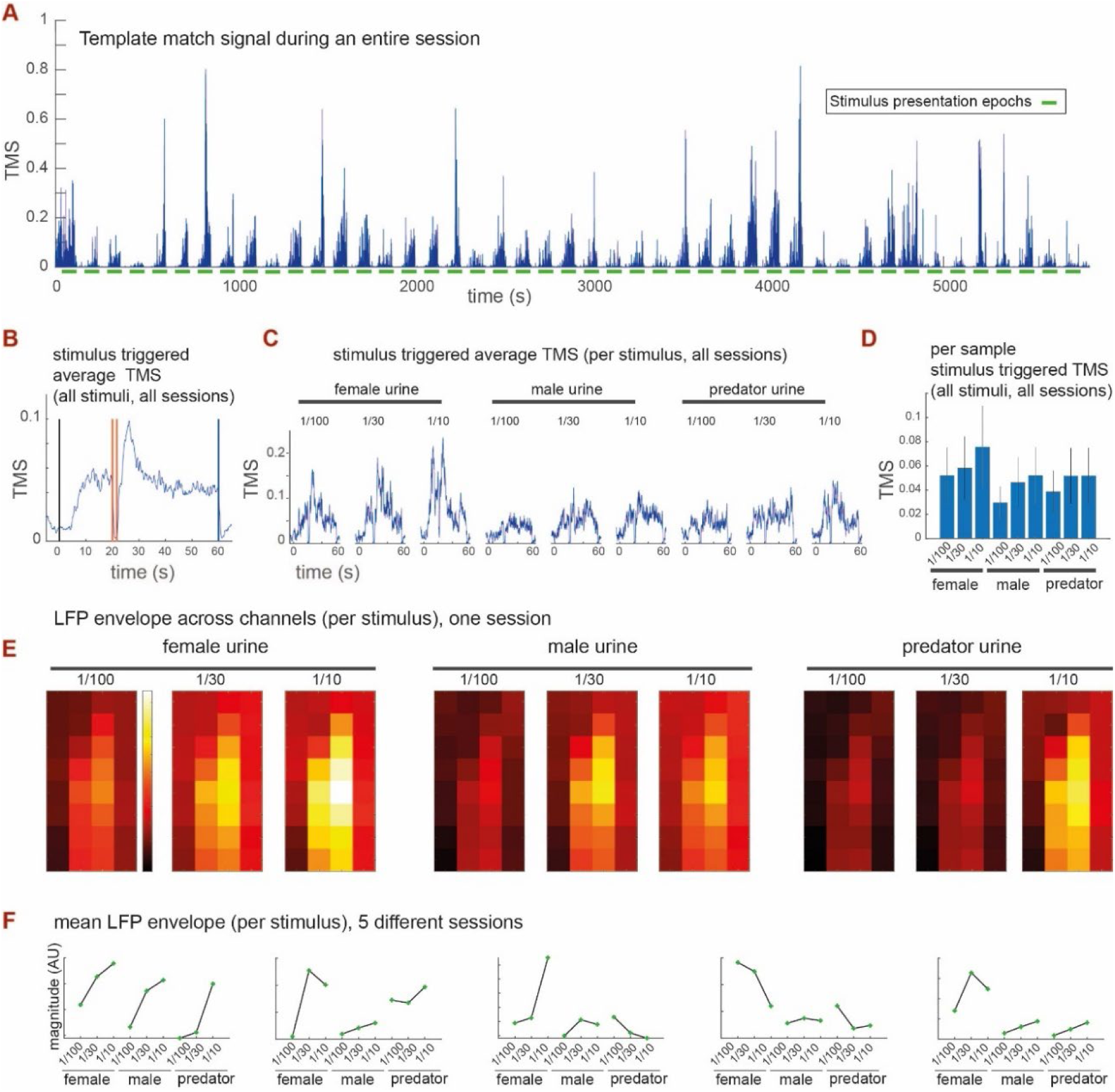
Stimulus dependence of the TMS and the LFP envelope. **A.** An example of the TMS during an entire recording session. Horizontal green lines correspond to stimulus presentation periods (60 s), starting with application to the nostril, and ending 40 seconds after nerve stimulation. Note how the TMS increases during stimulus presentation periods. **B.** Stimulus-triggered average TMS (staTMS), around all stimulus presentation times, in one type of experiment (D2, **Table S1**). Note that the signal rises following stimulus *application* (time 0), and then again during electric *stimulation*. The black vertical line corresponds to stimulus *application*. The two red vertical bars correspond to the start and end of the 1.6 s nerve stimulation. The blue vertical line marks the beginning of the wash period, which is effectively the end of the stimulation period. During electrical stimulation and washing, the TMS drops since electrodes record stimulation / flushing artefacts, respectively. **C.** Stimulus triggered average TMS (staTMS) for each stimulus. **D.** The mean template signal (directly proportional to the areas in C) and the standard error across all presentations of each of the stimuli. **E.** LFP envelope magnitudes associated with each of the stimuli. Note the stimulus and dilution dependence of the signal. **F.** Mean LFP envelope across all sessions in this type of experiment (D2, **Table S1**). Each panel shows values for one session. The leftmost panel corresponds to the data in E.

### Both the TMS and the LFP envelope depend on stimulus identity

We have shown above that the TMS rises during stimulus presentation. But is the TMS merely a reflection of stimulus uptake, or does it instead depend on the chemical composition of the stimulus? The fact that TMS magnitude varies between presentations (**Fig. 5A**), already suggests that the network pattern reflected by it might vary depending on the stimulus. To address this hypothesis, we derived the staTMS separately for each stimulus (averaged across all sessions with the same stimulus set). Comparison of these staTMSs reveals a stimulus-specific and dose-dependent response, with more concentrated stimuli eliciting higher staTMS profiles (**Fig. 5C**). Larger responses to female over male stimuli are also consistent with previous reports by us and others (5, 6, 67). Comparison of TMS distributions across samples and sessions, averaged for each stimulus, yields highly significant differences (**Fig. 5D**). Specifically, when comparing TMS value distributions (Kruskal-Wallis test, across all samples, treated as independent samples), the p-value for a main effect of stimulus is essentially 0 (see Methods). Similar conclusions regarding stimulus dependence are reached when applying the same analysis to another stimulus set (**Fig. S7B, C**).

Recall that the TMS is merely a measure of *similarity* to the ONED. Thus, high TMS values do not necessarily reflect high LFP intensities. To test if LFP *intensity* is also stimulus-dependent, we derived the average LFP envelope. Here, we consider time points during which the TMS is above threshold (see Methods, **Fig. S3C, D**). An example of mean LFP envelopes in one session, for each of the nine stimuli, plotted according to spatial location, is shown in **Fig. 5E**. This example shows that, like the TMS, the LFP magnitude (envelope) is also higher during presentation of less diluted stimuli. The mean LFP envelopes (averaged across the 32 channels) for all five sessions from this data set are shown in **Fig. 5F**. These examples demonstrate that, in all sessions, the LFP magnitude varies with stimulus identity, and in many of them, with stimulus dilution as well.

Taken together, our results indicate that both the TMS, a measure of similarity to the LFP distribution during oscillatory events, and the LFP distribution, are stimulus dependent. This suggests that more “intense” stimuli elicit higher LFP fluctuations, with full-blown oscillatory episodes representing an above-threshold state of a continuous activity pattern reflected by the TMS.

### Relationship between LFP oscillations and single unit activity

We have shown that the oscillatory network state, as reflected by the TMS, is related to stimulus delivery. Because AOB single-unit activity typically increases following VNS stimulus presentation (4, 5, 9, 68–70), the TMS and spiking rates should be correlated. However, what is the precise relationship between these measures? The TMS and the summed rate (combined across all recorded single and multi-units) during an entire session are shown in **Fig. 6A**. Expectedly, they appear correlated at this coarse time scale. Yet, inspection of the relationship at a finer temporal resolution reveals that the overlap is not precise (**Fig. 6A** bottom).

**Fig. 6.**
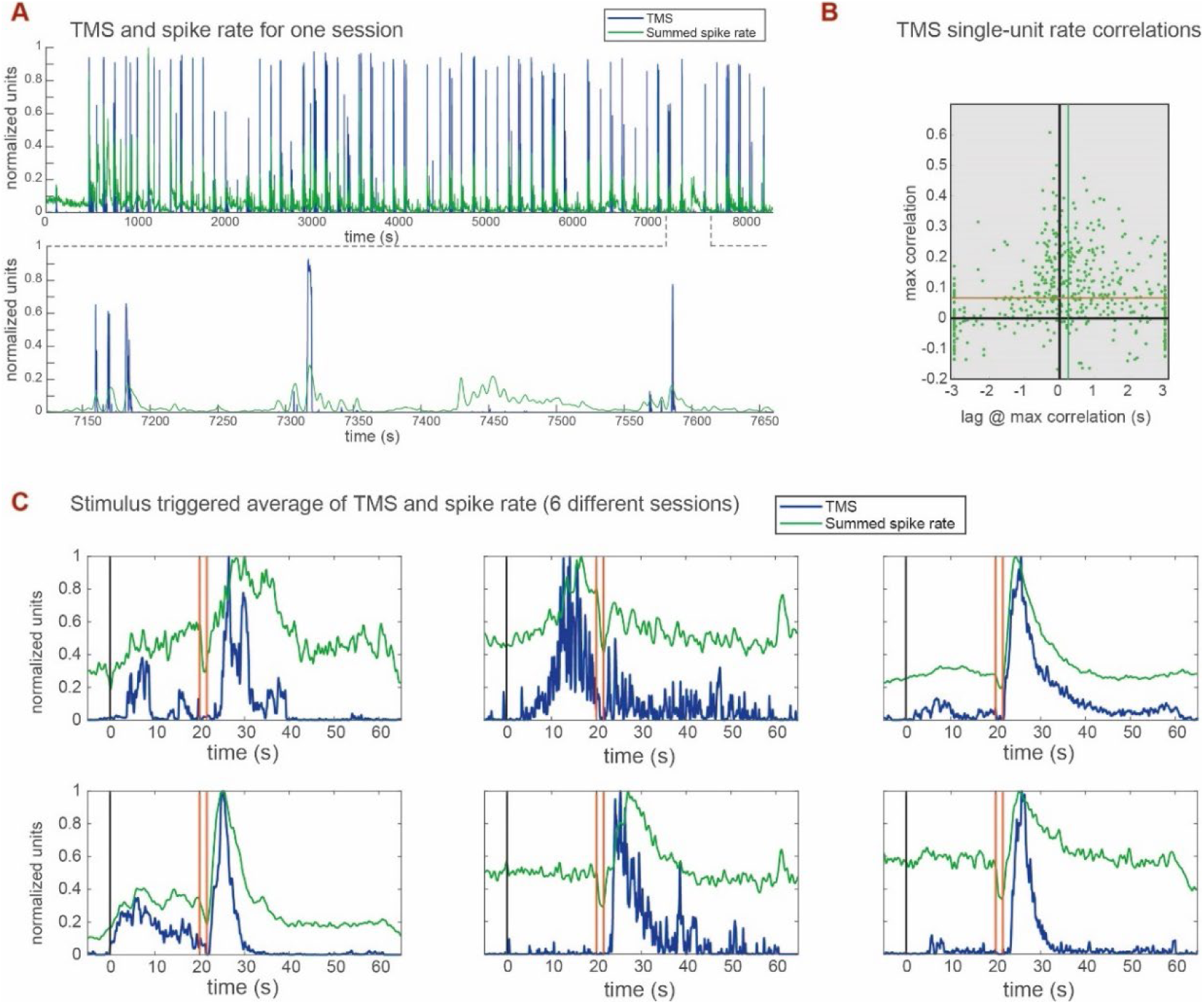
Relationship between TMS and unit activity. **A.** LFP template match (blue) and summed neuronal activity signal (green) during an entire session. A temporally zoomed view is shown below. Although both signals are correlated by virtue of their stimulus dependence (correlation of 0.565 at a lag of 0.076 s), closer examination shows that they do not perfectly overlap. Both single and multi-unit data are included in this panel. **B**. Maximal cross covariance magnitudes and lags at which they were obtained. Each dot represents one single unit (n = 520). Usually, the highest CC is achieved when unit activity follows the TMS. The medians of the distributions of CCs (red lines) and lags (green lines) are both positive. **C**. Stimulus triggered averages of the TMS and the summed single unit activity (PSTH). Averages of both measures were calculated across all stimuli in each session. Each of the six panels represents one session. See **Fig. S8** for more examples.

To reveal the temporal relationship between the two signals, we calculated the maximum cross-covariance (CC) between the firing rate of each single-unit (n = 520) and the TMS corresponding to that session, and the lag at which it was obtained (**Fig. 6B**, a positive lag indicates that the TMS precedes the single-unit rate vector). Most, but not all of the units are positively correlated with the TMS. While lags associated with high CCs are either positive or negative, the highest correlations occur for smaller lags. Across all units, the median CC is 0.067 (sign rank p-value: 1.03•10^-42^) while the median lag is 0.247 s (sign rank p-value: 0.002).

To explore the relationship between these signals in the context of sensory processing, we superimposed, for each session, the staTMS and the single-unit-summed peri-stimulus time histogram (summed-PSTH). Examples of this comparison for six sessions are shown in **Fig. 6C** (see also **Fig. S8**). Although these plots suggest that the TMS has a sharper onset and offset compared to the summed-PSTH, TMS derivation includes a sharpening (contrast-enhancing) step, limiting the validity of this comparison. However, this sharpening does not affect the peak time, and it is apparent that in many cases, both signals peak together, strongly suggesting that they are physiologically related. Notably (see also **Fig. S8**), the average TMS generally provides a more reliable indication of sensory stimulation than the combined spike rate. This is because single-unit sampling is sparser and random, with marked differences in yield of stimulus responsive neurons across sessions. These observations not only highlight the relevance of the TMS for sensory processing, but also as a simple and reliable (yet low dimensional) indicator of stimulus-induced VNS activity.

### Oscillations are most prominent in the ECL and likely reflect a reciprocal exchange between AOB-MCs and GCs

Having shown the relationship between oscillatory events and single unit activity, and their dependence on stimulus presentation, we turn to investigate their network source. Most of our recordings (see **Table S1**) were conducted with the planar electrode array advanced *in parallel* to the AOB ECL (**Fig. 7A**), also denoted as the mitral cell layer, or external plexiform layer (10). The ECL contains somata of AOB-MCs and displays characteristic signatures of spiking activity (4, 5). Although we dip electrodes in a fluorescent dye to routinely confirm within-ECL location *post-hoc*, reconstructions are not accurate enough to assign individual contact sites to specific layers. Yet, we can gain important information about electrode localization by comparing hotspots of single-unit neuronal activity with LFP energy distributions. Comparison of LFP signatures and single unit activity in individual sessions reveals a clear correspondence between oscillatory LFP power and neuronal firing activity, which is often associated with stimulus-induced neuronal responses (not shown). Such activity patterns are commonly associated with the ECL, highlighting the correspondence between LFP signals and AOB-MC activity. An example from one session showing the LFP signature and single unit activity is shown in (**Fig. 7C, F**).

**Fig. 7.**
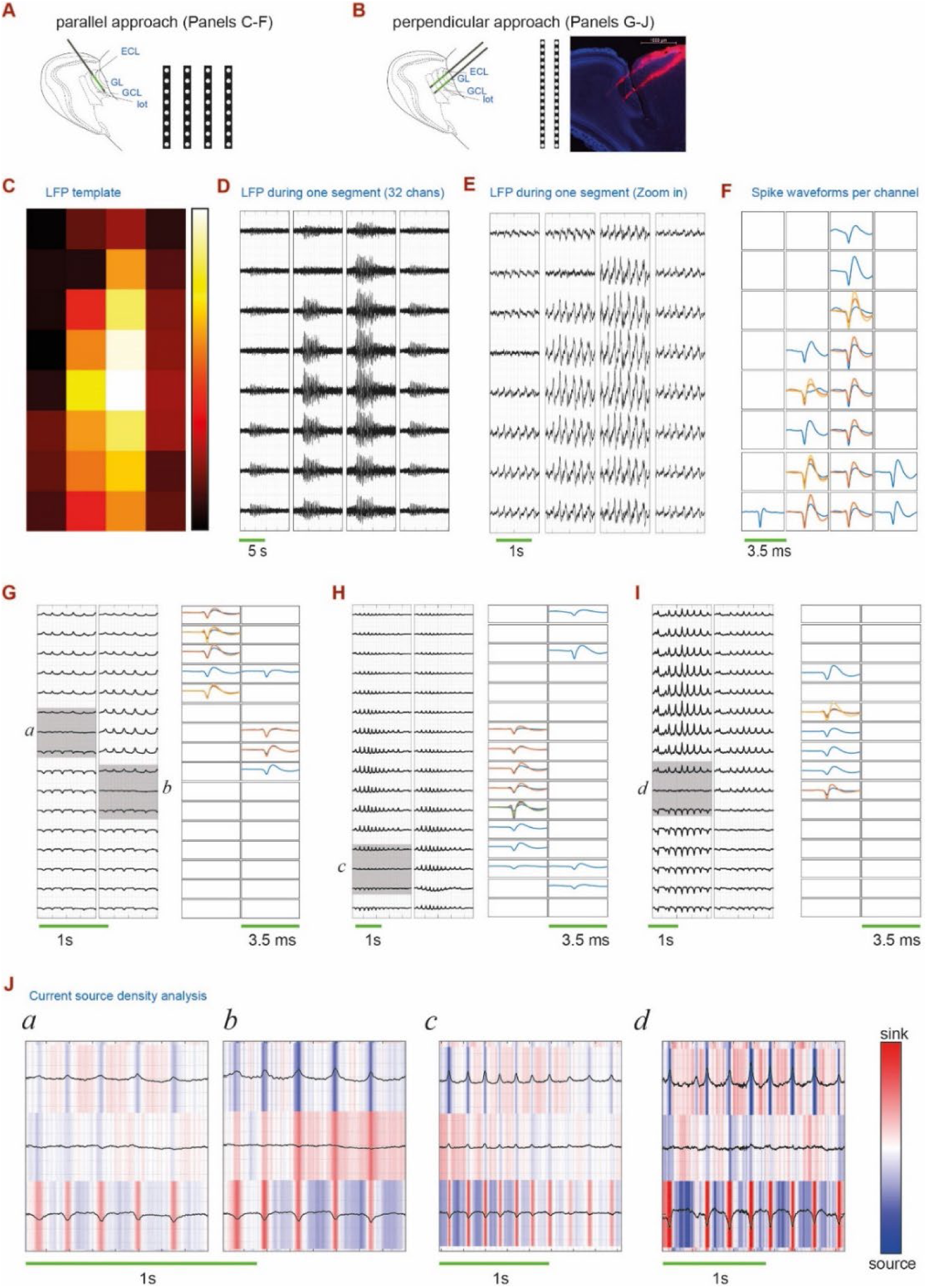
Evidence for localization of oscillations within the AOB ECL. **A.** Schematic of a parallel penetration into the AOB. **B**. Schematic of a perpendicular penetration into the AOB, using probes with a 2×16 configuration. Also shown is a dye tract of the electrodes in the session shown in panel G. **C.** Spatial spread of LFP energies in one of the sessions (also shown in Fig. 1). **D.** LFP oscillations during one oscillatory episode, across all channels, plotted according to spatial position. **E.** Magnified temporal view of D, highlighting the phase similarity across all channels. **F**. Spike waveforms for the sites shown in C. Each square shows the spike waveforms recorded in that channel. Both single unit and multi-unit activity waveforms are shown. Each trace in each panel corresponds to one single-unit or multi-unit. Although a given neuron can be recorded by multiple channels, here, we plot the waveform on the channel in which it had the largest amplitude. Note the correspondence between spiking activity and LFP amplitudes. **G.** Left - LFP signals during one episode recorded with a 2×16 electrode array. Note the reversal of the LFP along the probe (shaded regions). Right - **s**pike waveforms recorded on the sites shown on the left. **H, J**, like G, but for episodes from two different sessions. Note that in G, H, I that spiking activity is confined to sites at or above the LFP reversal. **J**. CSD analysis of the areas indicated by gray squares in G, H, and I. The relevant sections are indicated by the italicized letters. These CSD plots illustrate periods with reciprocal change between sinks and sources around the reversal point. Temporal scales in all plots are indicated by the green horizontal bars.

If LFP waveforms measured during parallel penetrations originate from the same neuronal structure, the oscillations are likely to be in-phase, without reversal of polarity. **Fig. 7D** shows all LFP channels during one oscillatory episode. The same episode is shown in expanded time view in **Fig. 7E**. This representative example shows that all channels within a session are indeed in-phase, suggesting that they originate from a homogenous neuronal structure (see **Fig. S9** for another example).

Continuing this line of reasoning, if oscillations result from an exchange between neuronal elements in different AOB layers, then their polarity should reverse at the boundary between these layers. To test this prediction, in a subset of experiments (see **Table S1**), we advanced electrodes *perpendicular* to the AOB, spanning its layers (**Fig. 7B**). Indeed, in most of these penetrations, we observe a clear LFP reversal between deeper and superficial sites. Because this is a crucial point, we show examples from three different sessions (**Fig. 7G-I**). The simplest interpretation of this reversal is that it represents the transition along the lateral olfactory tract, which separates the ECL from the AOB granule cell layer (GCL). Consistent with the idea that the AOB GCL activity is seldom detected by extracellular recordings, we observe neuronal spiking activity above, but not below the reversal (compare the left and right images within each panel in **Fig. 7G-I**).

Further support for our hypothesis that LFP reversals represent the transition between ECL and GCL is gained from current source density (CSD) analysis. CSD is the second derivative of the electric potential with respect to position, and provides a measure of current sources and sinks (71). We apply this analysis along the length of the recording probes that contain equally spaced recordings sites (and ignore the spread between adjacent electrodes). **Fig. 7J** shows the CSDs (represented by colors) along the sites of LFP polarity reversals (the shaded regions in **Fig. 7G-I**), superimposed on the original LFP traces. The CSD plots reveal an exchange of sinks and sources within each cycle period. Importantly, the CSD itself is reversed, in a manner matching the LFP reversal. As detailed in the Discussion, this interpretation is further supported by previous descriptions of LFPs and CSDs in AOB slices (72).

Integrating our observations, we propose the following scenario: VNO activation elicits spiking activity of single AOB neurons. With sufficient single unit activation, the network begins to recruit GCs, which supply inhibitory feedback to AOB-MCs and entrain oscillations. These oscillations then persist as long as incoming inputs provide sufficient drive to AOB-MCs. Once AOB-MC firing rates drop (due to reduction of incoming inputs), reduced GC activity ceases to drive oscillations. Under this scenario, full-blown LFP oscillations represent the peak of stimulus-induced AOB activity. This is consistent with the observation that oscillatory episodes (see **Fig. 3A, C** and **Fig. S4)** are briefer than response durations of many individual AOB single units (73).

### The effects of phase-dependent firing markedly increase with neuronal population size

A key hypothesis that emerges from our work is that neuronal oscillations play a role in information processing by modulating the timing of AOB neurons. For that to occur, the degree of phase locking observed here (e.g., **Fig. 4**) should suffice to induce a substantial effect on downstream neurons. We now present a simple calculation to demonstrate that even modest deviations of individual neurons from uniformity add up, and become extremely meaningful when considering co-active neuronal *populations*. Adopting the perspective of a downstream neuron receiving a fixed number of inputs, we calculate the probability to obtain a particular number of spikes within a specific integration window. We compare this probability between a uniform and a non-uniform distribution of firing within a cycle, keeping the overall firing rates constant. For example, we can compare neurons that fire at a constant rate of 5 Hz throughout the cycle, to neurons that fire at 4.5 Hz within one half of a cycle, and at 5.5 Hz during the other half (i.e., during the negative phase of the LFP). We then calculate the probability to obtain a particular number of spikes within a given part of the cycle (e.g., the negative phase of the LFP).

Specifically, we ask how the probability to obtain a given number of action potentials within a window depends on the firing rate of each neuron within the window, and the population size. This probability can be calculated using the binomial cumulative density function (Methods). As shown in **Fig. 8**, the differences, which increase with population size and the extent of phase locking, can be profound. For example, consider a population of 50 neurons, an integration window of 100 ms (half of a 5Hz theta cycle), and a baseline probability of 5 Hz (uniform firing), compared to a mildly locked non-uniform distribution (50% increased probability = 7.5 Hz within the window). Under these conditions, the probability to observe 30 spikes within that window is 0.14 under the uniform distribution, and 0.88 with the phase locked distribution. Thus, under these conditions, a mild degree of phase locking can increase the probability of observing 30 spikes within a 100 ms window six-fold! Larger effects are attained with stronger phase locking and larger populations (**Fig. 8**). In conclusion, a modest degree of phase locking, as observed here, can exert dramatic effects on downstream neurons with sufficiently large populations.

**Fig. 8.**
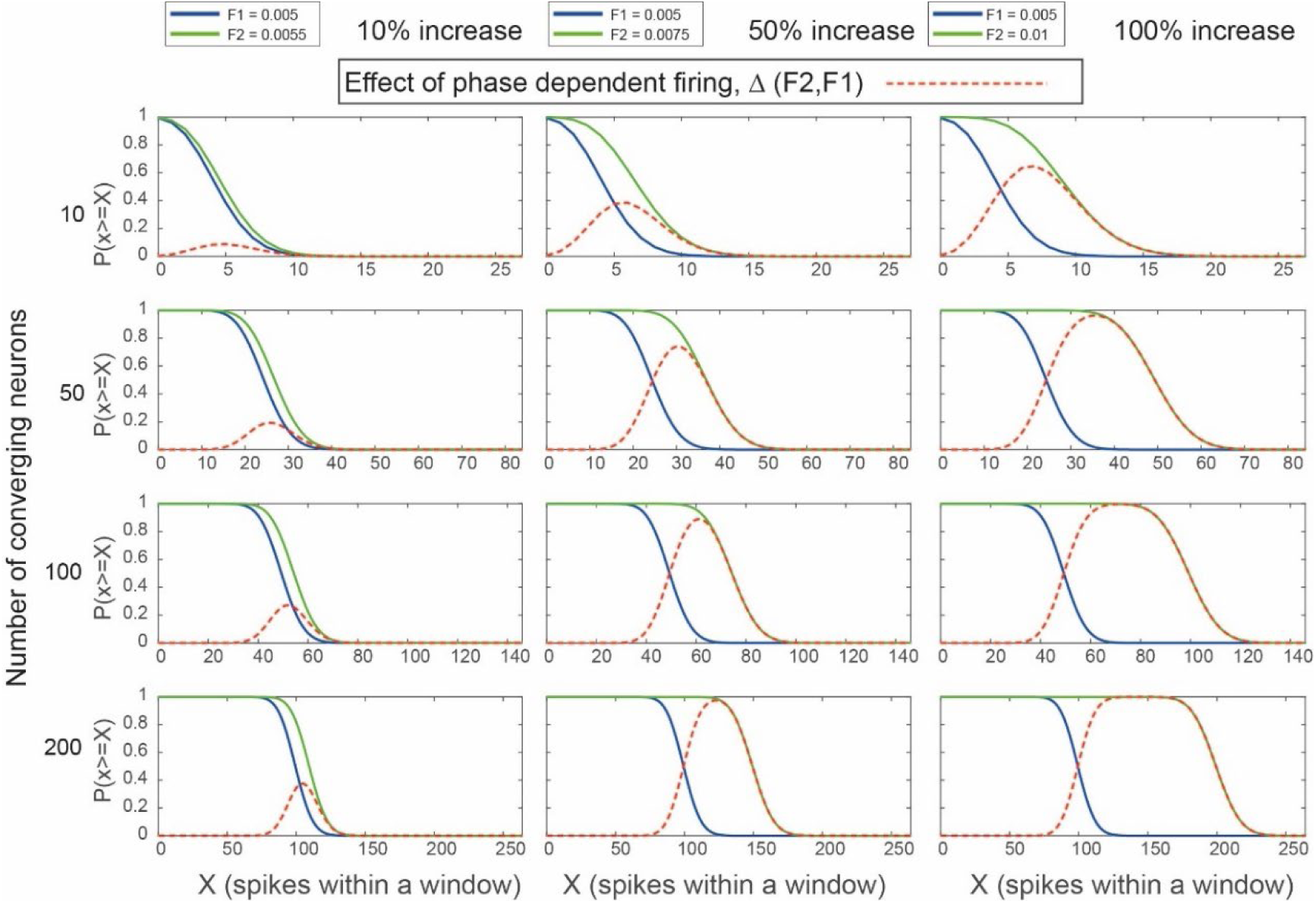
Effects of phase dependent firing on the probability of coincident spikes. Each panel shows the probability to obtain a certain number of spikes (or more), within a single integration window of 100 ms (i.e., half a 5 Hz theta cycle). Each row represents a different number of neurons (from top to bottom: 10, 50, 100, 200) and each column represents a different degree of spike time locking, quantified by the firing rate within that period. In all of these examples, the baseline rate (without spike time locking) is 5Hz which corresponds to a probability of firing of 0.005 within a 1 ms window. In each column, the degree of locking is different, and increases from left to right, assuming rates that are 1.1x, 1.5x, or 2x the baseline rate. Note that a 1.5-fold increase implies that 60% of all spikes are generated within half a cycle while a 2-fold increase within half a cycle from a uniform distribution implies that 2/3 of the spikes are emitted within half a cycle. In all plots, blue traces indicate the probability under uniform firing, and green traces indicate the probability with phase dependent firing. Broken red traces represent the difference between the two cumulative probabilities. This analysis highlights the benefit of even modest spike phase locking. The difference between uniform and phase dependent distributions increases with the number of neurons and the degree of locking (and remains true across all realistic baseline firing rates). Note the different horizontal scaling between the different rows.

## Discussion

In this manuscript, we studied neuronal mechanisms underlying processing of social information in mice. Specifically, we focused on oscillatory LFP patterns in the AOB. While spiking activity is arguably the most informative measure of neuronal communication, LFPs can reveal population level processes that are not easily observed by the activity of (even many) single units. We provided evidence that LFP oscillations are generated locally within the AOB, are associated with stimulus sampling, and occur even in the absence of arousal or active investigation. We have shown that they fall within the theta (2-12 Hz) band, but their sporadic appearance indicates that they are distinct from breathing-associated theta band MOB oscillations. Most importantly, we demonstrated that these oscillations exert a marked influence on neuronal spike timing, which can profoundly influence downstream processing. We thus propose that these oscillations shape AOB outputs as they are relayed to pathways that control behavioral and physiological function.

### The template matching approach

We discovered these sporadic theta oscillations by manually browsing LFP data. While often they were obvious, sometimes they were weak and obscured by noise associated with various mechanical and electrical events that our experiments involve. These intrinsic constraints impeded a systematic analysis of their relationship to stimulus presentation and unit activity. To overcome this challenge, we developed a semi-automatic method to identify oscillations (**Fig. S3**). Our approach relies on the highly conserved nature of envelope distributions within a recording session (**Figs. 2C, Fig. S1**). Using the mean energy distribution as a template (the oscillating network energy distribution, ONED), we scanned the entire recording session, thereby generating the continuous TMS. Initially, we expected the TMS to capture bona-fide oscillatory episodes. However, we found that it typically waxed and waned during the entire session, with a clear correlation to periods of stimulus presentation, and to stimulus identity (**Fig. 5, Fig. S7**). Thus, the TMS provides a measure of stimulus related network activity, with peaks representing full-blown oscillations following efficient recruitment of the AOB network. On a practical note, the TMS, which is sensitive to the spatial pattern of activity, provides a reliable way to monitor stimulus-induced activity in the AOB and, potentially, other regions as well. LFP signals reflect the activity of many neurons and are considerably easier to record than single-unit activity and, thus, a small number of electrodes sampling LFP activity in the AOB can provide a reliable readout of sensory activation.

### Source of the oscillations

The reversal of the LFP signal, and its comparison with the spread of unit activity (**Fig. 7**) strongly suggest that the origin of the oscillations is the AOB ECL. In addition, important insights can be gained from previous work in AOB slices. Specifically, Jia et al. (72) recorded LFP signals in different AOB layers during vomeronasal nerve activation. Based on LFP waveforms and CSD analysis, stimulation-induced LFP signals were divided into three periods. The first, strongest in the vomeronasal nerve and glomerular layers, was attributed to activation of vomeronasal sensory neuron axon terminals. The second period included a current sink in the glomerular layer and a source in the ECL. This sink/source pair was attributed to depolarization of apical AOB-MC dendrites. In the third period, a large sink appeared in the ECL with a corresponding source in the GCL. The ECL sink was attributed to GC peripheral dendrite depolarization, which can be induced by synaptic activation of AOB-MCs. Using whole-cell recordings from AOB-MCs and GCs, Jia et al. observed that the second LFP period coincided with excitatory post synaptic potentials in AOB-MCs, which evolved into action potentials. In the third period, inhibitory post synaptic potentials appeared in AOB-MCs, along with excitatory potentials in GCs. This inhibition was associated with reduced AOB-MC action potential generation. The three phases correspond to sequential activation of three main AOB circuit elements: vomeronasal nerve layer, AOB-MCs (spanning the glomerular layer and ECL), and GCs (spanning the ECL and GCL). Based on these analyses, and our observations, we propose that the oscillations observed here result from ongoing AOB activation, which, if sufficiently strong, leads to reciprocal exchange between AOB-MCs and GCs. We speculate that stronger sensory activation not only increases the rate of AOB-MC activation, but also that of GCs, resulting in increased inhibitory drive onto AOB-MCs and hence higher oscillatory frequencies. As the incoming signal wanes, there is no longer sufficient drive to activate AOB-MCs and GCs in synchrony, and oscillations decelerate and, eventually, disappear. Thus, frequency changes within a single oscillatory episode may result from dynamic changes in AOB-MCs inputs. It is important to note that, while these AOB oscillations fall within the theta band, they are mechanistically distinct from MOB theta oscillations (28). Since all our experiments were conducted in anesthetized mice, and most of them included tracheotomy, we conclude that the oscillations are triggered by sensory activation and are not dependent on active exploration or feedback from other areas. In a few experiments (see **Table S1**), a tracheotomy was not performed (and there was no flushing of the nasal cavity). In these experiments too, we did not observe ongoing breathing-associated theta oscillations, further supporting the notion that breathing does not play a direct role in the oscillations studied here.

### Previous studies on AOB theta oscillations

Our study of theta band LFP oscillations in the AOB follows previous reports. In addition to the *in vitro* analysis mentioned above (72), work from the Brennan (58, 59) and Lanuza laboratories (57) revealed stimulus-induced theta band oscillations in the AOB (see Figure 1 in (59) and Figure 2 in (74)). The former study also showed that oscillatory LFP bands can change following mating. Recordings in rats (61) revealed that coherence among theta LFP in various limbic regions (including AOB) depends on behavioral context. The widespread distribution of theta oscillations throughout limbic regions suggests that the patterns observed here may induce, or resonate with downstream VNS regions. While our recordings indicate that oscillations can arise in the absence of arousal, the work of other groups highlights the relevance of oscillations in the freely moving state. Another prominent class of AOB oscillations, described by us and others, are infra-slow oscillations (53–55, 75). These are much slower than those observed here, and likely play a different role, and arise from distinct mechanisms. Nevertheless, there may be a direct relationship between neuronal ensembles such as identified in (53) and those proposes here (see below and **Fig. 9**).

**Fig. 9.**
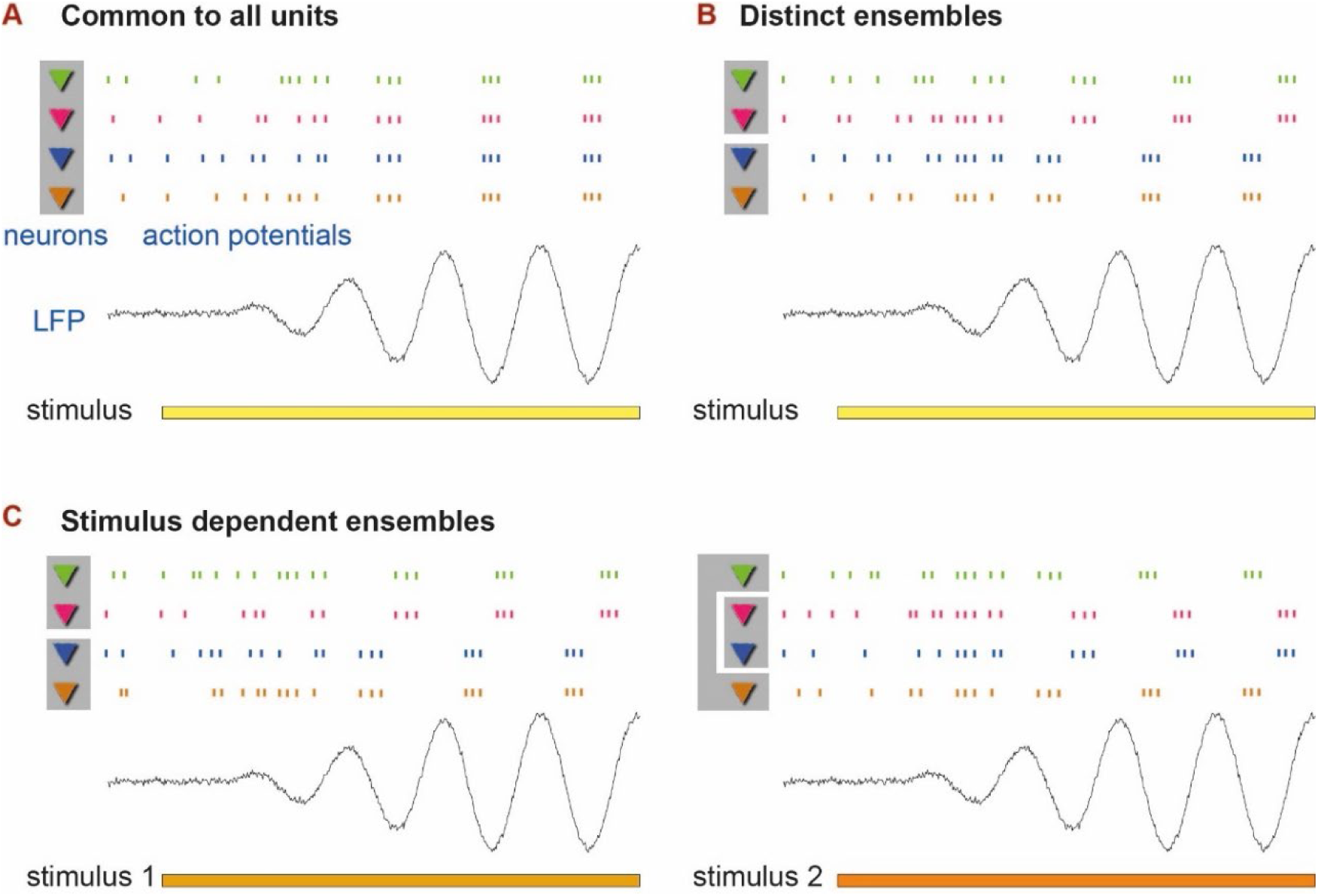
Models of physiological roles of theta oscillations and the effect of phase concentration. Cartoons of three different scenarios for phase dependent spike timing. Each scenario includes spike trains from four neurons, and a few LFP oscillations periods. Stimulus presentation is indicated by the colored horizontal bars. Shaded regions behind neuron icons represent co-active ensembles. **A.** Uniform locking of spikes from all neurons regardless of the stimulus. Here, oscillations form a common gating and amplification mechanism for all units and sensory stimuli. **B**. Locking of spikes, so that each unit has a preferred phase within the broad region of preference. Here, there may be distinct ensembles that represent distinct stimuli. **C**. Stimulus and unit dependent phase locking. In this scheme, an elaboration of B, ensemble memberships may change as a function of the stimulus.

### Mechanisms underlying generation of the oscillations

Mechanistically, we proposed that AOB theta oscillations result from an interaction between AOB-MCs and GCs. This conclusion is based on spatial analysis of LFP waveforms (**Fig. 7**) and previous work in slices (72). Notably, however, in the MOB these circuit elements were proposed to underlie the much faster gamma rhythm (29, 30, 34, 76). This apparent discrepancy may be reconciled by the fact that AOB and MOB neurons differ in intrinsic properties (48, 49), connectivity (10), and the inputs they receive. All of these features (and others) can markedly affect oscillation frequencies (29–31) and potentially account for the phenomenology of the oscillations observed here. Further experimental and modeling studies of the AOB network are required to pinpoint the mechanisms that produce AOB oscillations.

The role of active sampling and its influence on sensory processing is widely discussed in the context of the MOS, where sampling patterns can modulate the activity of sensory neurons, and hence of downstream targets (43, 44, 77–80). This suggests another potential source for the oscillations, namely, the vomeronasal pump. While there is evidence for intrinsic rhythmicity of the pump (46, 81), it is not known if it plays a role in the oscillations that we see here. One can speculate that more intense investigation will give rise to different pump dynamics, and hence alter sensory neuron activity patterns, in turn shaping AOB input patterns. To reveal the effect, if any, of active sampling on these oscillations, it will be necessary to record them in awake behaving animals. Alternatively, VNO sampling may not be as amenable to controlled voluntary modulation as is sniffing, raising the idea that the AOB realizes an intrinsic oscillatory mechanism serving a similar functional role (i.e., gating, binding, and multiplexing). Along the same lines, inputs from other brain regions could modify AOB neuron excitability and thus modulate the presence and expression of oscillations. This possibility can be investigated by enhancing, or diminishing, feedback inputs from other brain regions (82–86).

### The physiological relevance of the oscillations

The existence and prevalence of the stimulus-induced theta oscillations beg the question whether they play a role in information processing. We strongly advocate the view that they are not an epiphenomenon, but rather reflect an important aspect of AOB physiology. Below we propose several models, all of which are inspired by previous work in the MOS (23, 27, 31, 32, 39, 64, 76, 87-95), and other regions (26, 96).

One prominent aspect of the VNS and the AOB is its integrative structure. Along with established patterns of labeled line coding in the AOB (13, 16, 97), many natural stimuli are complex blends of molecules (5, 8, 17), and this implies that stimulus recognition may involve integration of information across channels (98). Indeed, AOB circuitry itself already realizes substantial integration, as AOB-MCs often sample information from multiple glomeruli (1, 10, 11, 51). If distinct channels convey related information, there may be an advantage for these channels to be co-active. Thus, one simple hypothesis (**Fig. 9A**) is that the oscillations serve to constrain neuronal firing to limited temporal windows, and thus activate downstream regions more efficiently. In this scenario, oscillations serve a gating and amplification function: low neuronal activity levels will not give rise to oscillations, but once the activity is high enough, oscillations will appear, constrain spike timing to smaller windows, and facilitate information transfer. This scenario is supported by the results shown in **Fig. 4B, F, G** and the analysis in **Fig. 8**. We stress that this model, as well as the others proposed below, rely on the observation that LFP oscillations are in phase throughout the extent of the AOB ECL (**Fig. S6**).

A more elaborate hypothesis suggests the existence of multiple ensembles, so that members of each are activated together. Under this scenario (**Fig. 9B**), neurons within an ensemble share a similar phase preference and are thus more likely to exert a synergistic effect on downstream targets. The notion that different units follow distinct phase preferences is supported by the analysis shown in **Fig. 4C**. However, unlike the MOS, where spike times of neurons “tile” the theta breathing cycle (40, 64), here most of the activity is concentrated within about a third of a period. Finally, the most elaborate scenario holds that phase distributions of units are stimulus-dependent (**Fig. 9C**). According to this model, each neuron can participate in different ensembles depending on the stimulus. Support for this idea comes from the analyses shown in **Fig. 4D, E**. The two latter models may seem inconsistent with the idea, promoted also by us (73) and others (49), that the VNS is slow and temporally imprecise. However, there is no discrepancy between our previous conclusions and the present observations, since the synchronization proposed here involves joint activity of neurons, rather than their locking to sensory stimulation on a trial-by-trial basis.

### Potential effects of coordinated activity on downstream processing stages

As we have shown above (**Fig. 8**), even a modest degree of phase-dependent firing can exert a profound effect on joint neuronal firing probability within a limited temporal window. The actual effect depends on the integrative properties of downstream processing stages, whose details are not known to us. This includes the degree of anatomical convergence, and the spatial and temporal synaptic integrative properties of neurons in these regions. Presently, our findings highlight and implicate a potential mechanism that cannot only gate, but also bind and organize activity from distinct neuronal elements (**Fig. 9**). Understanding which, if any, of these mechanisms play a role in AOB information readout remains a central goal for future research.

## Materials and Methods

### Subjects

For recordings, adult sexually experienced male and female mice from the BALB/C and ICR strains were purchased from Envigo Laboratories (Israel). All experiments were performed in compliance with the Hebrew University Animal Care and Use Committee. A detailed description of the strains and sexes used in each session appears in **Table S1**.

### Stimuli

Stimuli were collected from adult estrus and non-estrus sexually-naïve female mice of the BALB/C, C57BL/6 and ICR strains (Envigo Laboratories, Israel), as well as castrated, naïve, and sexually experienced male mice from the BALB/C, C57BL/6 and ICR strains. Some experiments also included a mixture of predator urine (purchased from PredatorPee, Maine, UAS) or a ringer’s control stimulus. For mouse urine collection, mice were gently held over a plastic sheet until they urinated. The urine was transferred to a plastic tube with a micropipette and then flash-frozen in liquid nitrogen and subsequently stored at -80°C. Three of the sessions were previously analyzed in a different context and include urine, saliva, and vaginal secretion stimuli, as described in (8). One session used urine stimuli with different pH values, and was previously included in (99). See **Table S1** for a detailed description of the stimulus sets.

### Experimental design

Experimental procedures were detailed in (62), and are described briefly here, with the differences noted. Mice were anesthetized with 100mg/kg ketamine and 10mg/kg xylazine, a tracheotomy was performed with a polyethylene tube to allow breathing during flushing of the nasal cavity, and a cuff electrode was placed around the sympathetic nerve trunk with the carotid serving as a scaffold. Incisions were closed with Vetbond (3M) glue and the mouse was placed in a custom-built stereotaxic apparatus where anesthesia was maintained throughout the entire experiment with 0.5-1% isoflurane in oxygen. In parallel penetrations, a craniotomy was made immediately rostral to the rhinal sinus, the dura was removed around the penetration site, and electrodes were advanced into the AOB at an angle of ∼30° with an electronic micromanipulator (MP-285; Sutter Instruments, Novato, CA). In perpendicular recordings, the craniotomy was made caudal to the rhinal sulcus and the electrodes advanced approximately at 60°. All recordings were made with 32 channel probes with either 8 channels on each of 4 shanks (NeuroNexus Technologies, Ann Arbor, Michigan) or 16 channels on each of 2 probes (see **Table S1**). Due to an oversight, some of the sessions include LFP recordings from only 16 channels (rather than 32, see **Table S1**). Unless indicated otherwise, data from these sessions are also included in our analysis, as the 16 channels (spanning the superficial half of the probe) were sufficient to derive the TMS (see **Fig. S2**). Before recordings, electrodes were dipped in fluorescent dye (DiI, Invitrogen, Carlsbad, CA) to allow subsequent confirmation of electrode placement within the AOB ECL, which contains the mitral-tufted cells (10). In each session, stimuli were typically presented 5 times in a pseudorandom order. In each presentation, 2 µl of stimulus was applied directly into the nostril. After a delay of 20 s, a square-wave stimulation train (duration: 1.6 s, current: ±120 µA, frequency: 30 Hz), was delivered through the sympathetic nerve cuff electrode to induce VNO pumping and stimulus entry to the VNO lumen. Following a second delay of 40 s, the nasal cavity and VNO were flushed with 1-2ml of ringer’s solution which flowed from the nostril, into the nasal cavity, and sucked out from the nasopalatine duct via a solenoid-controlled suction tube. The cleansing procedure lasted 50 s and included sympathetic trunk stimulation to facilitate stimulus elimination from the VNO lumen.

### Electrophysiology

Neuronal data was recorded using an RZ2 processor, PZ2 preamplifier, and two RA16CH head-stage amplifiers (TDT, Alachua, FL). LFP data was collected at 508.63 Hz and band-pass filtered at 0.5 to 300 Hz (“EEG” setting on the TDT system). As described below, a notch filter was applied to remove 50Hz line noise. Single-unit and multi-unit activity were sampled at 24414 Hz, band-pass filtered (300-5000 Hz) and custom MATLAB (Mathworks, Natick, MA) programs were used to extract spike waveforms. Spikes were sorted automatically according to their projections on two principal components on 8 channels of each shank using KlustaKwik (100, 101) and then manually verified and adjusted using the Klusters program (101). Spike clusters were evaluated by consideration of their spike shapes, projections on principal component space (calculated for each session individually) and autocorrelation functions. A spike cluster was defined as a single unit if it had a distinct spike shape and was fully separated from both the origin (noise) and other clusters along at least one principal component projection, and if its inter-spike interval histogram demonstrated a clear trough around time 0 (of at least 10 ms). Clusters comprising more than one single-unit were designated as multi-units. Thus, using the present definitions, multi-units could represent the activity of as few as two units, or more. In most of the analyses here we used single-unit data. When using multi-unit data, this is indicated explicitly. We note that our multi-units are not equivalent to what is commonly referred to as multi-unit activity and which includes above threshold signals in each of the electrodes. The latter are not included in our analyses. As a further criterion of our single-unit data, we examined all spike shapes by calculating the shape symmetry. The symmetry is a good measure for accidental noise that was misclassified, which yields high symmetries. Using a graphical representation of all spike shapes, we examined the entire data set for such noise like spikes and these were excluded from all analysis.

### Data analysis

Our dataset contains recordings from 34 sessions (see **Table S1** for details). Generally, all analyses were conducted in MATLAB, using built-in MATLAB functions when available, or custom code as needed. Initial, exploratory data analysis was performed using a custom-written data browser that allowed viewing all LFP and electrode data, superposed with trial events and manual oscillatory event definition. The interface was instrumental for revealing the oscillatory episodes that form the basis of this study. Browsing the data, we occasionally observed distinct oscillatory events. In order to analyze these events, and their relation to other experimental and physiological variables, it was first necessary to find an objective way to detect them and define their start and end times. This turned out to be a challenge since these events (as we show in the manuscript) vary in their temporal and spectral characteristics. In addition, although their presence was often obvious (e.g., **Fig. 2A, B**), event onset and offset were typically gradual and thus difficult to delineate objectively. Finally, our experiments include inevitable sources of mechanical/electrical noise (associated with electric stimulation of the VNO, valve opening and closing, stimulus application, and rinsing of the nasal cavity) (62), which may overlap with oscillatory events in the temporal and/or the spectral domains. These considerations imply that a simple bandpass spectral filter will yield both false positives and missed detections, and will thus not allow effective event detection. To overcome these challenges, we exploited the fact that within a recording session, all events shared a similar across-channel energy distribution. That is, while envelope amplitudes during these events often differed in *absolute* values, their *relative* amplitudes across recording channels were highly similar (**Fig. S1**). This prompted us to use the average LFP envelope pattern across all manually defined events as a *template,* and apply it to search for similar activity patterns across the entire recording session. Specifically, we calculate the linear correlation coefficient between the template and the LFP envelope across all channels at each point in time. The result is smoothed, rectified, and sharpened to increase contrast, yielding a measure of similarity between ongoing network activity and the oscillatory event template (**Fig. S3A**). We denote this continuous measure as the *template match signal* (TMS).

Our data analysis pipeline includes 3 key stages:

Stage 1. Generation of the template. First, several oscillatory events were manually defined for each session (See **Fig. S1**). These were then used as seeds to derive the LFP template using custom code. The distribution of LFP energy across channels was then calculated for each of these events by calculating the mean LFP envelope, for each of the channels, across all samples. **Fig. 2C** shows these envelopes for each of the defined segments. The template itself (e.g., **Fig. 2D**) is simply the mean of these events’ envelopes. When averaging across manually defined events, we first derive the mean envelope associated with each event, and then average across events (so that each event has the same contribution, regardless of its duration). During this stage, we sometimes also identified and defined broken (and hence noisy) channels, which were excluded from TMS calculation. The template is then saved in a data file and used in subsequent processing stages. We stress that although there is an unavoidable level of subjectivity in this procedure due to the manual selection of the seed events, once defined, the episodes are derived automatically and objectively. Furthermore, the high stereotypy between all user defined events (e.g., **Fig. 2C, Fig. S1**) implies that the template, and hence the TMS, are minimally sensitive to the precise choice of manually defined events.

Stage 2. Analysis of the data in each session using a wrapper function. Given a data session as input, the function loads the LFP data, spike data (including both spike times and definitions of unit classifications, i.e., single vs. multi units), event data (e.g., trial events such as stimulus presentation and stimulation), the envelope template (from stage 1), and electrode configurations. Using this information, the function conducts multiple steps of analysis which are described below. The outputs are saved as MATLAB figures and a data structure (for each session).

Stage 3. Finally, we run several scripts that load the results data from multiple sessions, as generated by the previous stage, and conduct global analyses on this combined data. These scripts serve various purposes such as analysis of basic episode features, spatial distributions of the template, relationships between LFP and single-unit data, phase analyses and so on.

All data figures in this manuscript are generated by MATLAB, exported to adobe illustrator where they were resized or altered graphically (colors, line widths, etc.) for clearer presentation. *All the functions, scripts, data, and figure files generated by these functions are available upon request*.

Below is a description of the analysis steps included within stage 2, as applied to each of the sessions. Many of the calculations performed by the analyses scripts in stage 3 employ the same algorithms as in stage 2, and are described in the corresponding sections in the manuscript.

### Analysis procedure per session

1. Basic LFP processing. We load LFP data, apply a notch filter (to remove the 50Hz line noise) using the MATLAB *filfilt* function. The filter is defined with the *designfilt* MATLAB function using the following parameters: type: bandstopiir, FilterOrder: 20, HalfPowerFrequency1: 49, HalfPowerFrequency2: 51, SampleRate: 508.63. The LFP envelope is derived with the MATLAB *envelope* function with the *rms* option, and a window of 0.1 seconds.

2. Derive the LFP template match signature (TMS), by calculating the linear correlation (using the MATLAB *corr* function) between the template (derived in stage 1), and the LFP envelope at each point in time, across the active channels. The signal is then smoothed with a 0.5 s Gaussian window (using the MATLAB *smoothdata* function), rectified (so that negative values become 0), and raised to the 10^th^ power to enhance contrast (**Fig. S3A**). As described above and in the main text, the template is the mean LFP envelope across all channels (oscillating network energy distribution, ONED), in manually defined events. Note that low TMS values during electrical stimulation of the sympathetic nerve trunk and at the end of each trial (the start of wash period), result from the fact that during these stages, the spread of energies (induced by stimulation artifacts and other experimental sources of noise) is not correlated with that of the LFP energies during oscillatory episodes (see **Figs. 5A-C**, **Fig. S3**, **Fig. S7**).

3. Detect candidate episodes. We apply a threshold on the LFP template signal. We used as threshold the mean signal due to its simplicity, and the fact that it was typically lower than other thresholding methods that we tested. A lower threshold implies more detected episodes and hence larger samples for statistical analyses. After setting a threshold, we obtain a binary template signal. All such detected sections that are at least 1 s long are considered as potential episodes. An example of the TMS during an entire session is shown in **Fig. S3C**, with a time-expanded section in **Fig. S3D**. The bottom plots within these panels show the threshold and the binary TMS signal obtained after applying it.

4. Ridge extraction and phase derivation. Rationale: The spectro-temporal analyses in this work were based on the wavelet synchro-squeezed transform (WSST) (102, 103). We chose this approach because it includes a “reassignment” step that enhances temporal and spectral estimates by compensating for spreading effects of the base wavelet, and because it facilitates ridge extraction using the built-in MATLAB functions. A ridge is a sequence of (typically smoothly varying) frequencies and their amplitudes in time, and ridges are thus particularly suitable for analyzing signals that contain a significant chirp-like component, like the oscillations analyzed here. Multiple ridges can be extracted, with the first ridge being the maximum energy time-frequency ridge, and subsequent ridges showing decreasing energies. The inverse WSST (iWSST) reproduces the original signal in the time domain from the transform generated by the WSST. It is also possible to reconstruct the original signal using one or more ridges. If a ridge captures a substantial fraction of the energy of the original signal, then reconstruction based on it will yield a result similar to the original signal. In our analysis, we focused on episodes for which a single ridge can provide an above threshold reconstruction (*reconstruction score*, defined as the correlation coefficient between the original signal and its reconstruction). Analysis based on ridges rather than on a narrow frequency band is crucial because any given frequency band might either miss oscillatory components, or contain non-oscillatory components. In addition, instantaneous phases of signals that contain multiple frequency bands are often ambiguous, whereas signals that are composed of a single ridge contain a single frequency at each point in time and thus are associated with an unambiguous phase.

Practically, we consider the channel with the highest LFP energy and then apply the wavelet synchro-squeezed transform on this signal (using the MATLAB *wsst* function). Focusing on one channel is warranted given the high correlation at 0 lag among all channels in a given episode, as calculated by the cross covariance (MATLAB *xcov* function) of all channels (**Fig. S6**). We then remove all frequency contributions above 40Hz (which are substantially above our range of interest), and then extract the first three ridges using the *wsstridge* MATLAB function, using a *penalty* value of 5. Higher *penalty* values favor ridges without abrupt changes in the frequency domain. We then define the mean frequency of each ridge as the weighted mean frequency across all bins, where the weighting is given by the amplitude (the absolute value of the WSST in the corresponding bin). Next, we reconstruct the original episode using the inverse wavelet synchrosqueezed transform (*iwsst* MATLAB function). Episodes are reconstructed using the first ridge. We then calculate the correlation coefficient between the original signal and the reconstruction and derive the instantaneous phase of the reconstructed signal using the first ridge. The instantaneous phase is obtained by taking the angle (MATLAB *angle* function) of the Hilbert transform (MATLAB *hilbert* function). At this stage, we have a set of episodes, with associated phases, reconstruction scores, and other measures such as the mean frequency (as described above), mean template match and the mean envelope (energy).

5. Selection of episodes for further analysis. For further analysis, we selected only episodes that passed two criteria: a reconstruction score > 0.5, and a mean frequency of 2-14 Hz. The mean frequency is defined here as the mean across the first ridge, with the frequency in each time point weighted by the amplitude of the ridge during that sample. This range was selected after we conducted extensive manual inspection of oscillatory events and concluded that virtually all such oscillations are within the range. These two selection criteria were applied to exclude episodes without a dominant theta band component.

Next, we manually excluded episodes whose high reconstruction score is not due to a good reconstruction of the theta mode of the episode (“false positives”), and identified episodes that include obvious oscillatory modes, are noise free, and complete (i.e., that the entire oscillatory episode appears uninterrupted by noise) (“complete” episodes). The rest, which are the majority of episodes are designated as accepted episodes. Unless indicated otherwise, we used accepted and complete episodes in our further analyses. Complete episodes are mainly used for in-depth analyses such as CSD analyses. When a distinction is made between accepted and complete episodes, this is indicated in the text. Of 2609 candidate episodes, 266 were designated as false positives and thus excluded from further analysis, while 89 were designated as complete.

**Fig. S3 E-H** shows an example of the application of this procedure to one channel during one oscillatory episode. The original signal, and its reconstruction by applying the inverse-WSST, (iWSST) to the first ridge are shown in **Fig. S3E** in black and green, respectively. For this example, the reconstruction score is 0.84, indicating that the ridge provides a faithful account of the entire signal. The frequency (blue) and amplitude (green) of the first extracted ridge are shown in **Fig. S3F**, and the instantaneous phase is shown in **Fig. S3G**. **Fig. S3H** shows the original signal in the time domain (white), and the spectrogram of the first ridge. The mean frequency of the first ridge is 5.25 Hz, clearly within the conventional theta frequency range. **Fig. S4** shows the results of this analysis for all complete episodes, including reconstruction with the first, and for illustration, also with all first three ridges. Plots of the relationships between various features of each oscillating episode are shown in **Fig. S5A-D,** with complete episodes indicated by the red circles. In particular, **Fig. S5A** shows the relationship between the mean TMS and the reconstruction score. It can be seen that episodes with higher TMS values generally yield better first ridge reconstructions.

6. Current source density was calculated for each of the complete episodes as the second derivative of the LFP signal along the probes within a given shank (71). The CSD as a function of time is smoothed using the MATLAB *smoothdata* function with a gaussian window of 20 ms.

7. Analysis of the LFP and unit signals with respect to sensory events. To study the relationship between the TMS and the sensory events, we considered all stimulus presentation epochs. The stimulus presentation epoch begins with stimulus *application* to the nostril, and ends 40 seconds after the electrical nerve *stimulation*, for a total period of 60 seconds. Some of our analysis and displays include margins of 5 s at the beginning and ends of this epoch, for a total duration of 70 s per stimulus. For each of these epochs we derived the LFP template match signature, the binary template match signature (as defined above) and the spiking rates of each of the units (spiking rates were calculated in 0.2 s windows and then smoothed with the MATLAB *smoothdata* function, using a 1s *gaussian* window). With this approach we could retrieve the continuous TMS associated with individual trials, the mean template match signature associated with all trials of a given stimulus (within a recording session), and the number of samples in which the template match signature was above threshold. In addition, we derived the LFP envelope (that is the LFP rms signal itself rather than the TMS) associated with each stimulus. The LFP envelope was evaluated during samples in which the binary template signal was above threshold. Thus, the accuracy of the envelope estimation depends on the number of samples during which there was above threshold match with the template. In some cases, there were no such samples for a given stimulus and session.

8. To correlate firing rates of individual units (or all units together) with the LFP template signal, we first resampled the spiking rates by binning spike counts into bins that match the LFP sampling rate, and then smoothed them using a 5 s gaussian window (using the MATLAB *smoothdata* function). The relatively long window was chosen to reduce noise in the estimate of the firing rate and thus in the correlation between the spiking and LFP signals. The correlations between spike and LFP envelope signals were calculated using the MATLAB *xcov* function, with scaling set to *normalized*.

9. Spike time phase analysis. Spiking phases were only measured during oscillatory episodes (accepted or complete episodes as defined above). Recall that these are data episodes for which the first ridge mean frequency is within the theta range and that it yields a reconstructed signal with a correlation of at least 0.5 with the original full signal. Each sample in each episode is associated with an instantaneous phase (see above). Then, for each of the units, spikes that occurred within that episode are found, and for each we find the nearest LFP sample. This way, a phase is associated with each of the spikes of each of the units within each of the different stimuli (that were presented during that session). Importantly, during this process we also derive shuffled phase distributions. These are obtained by randomly reordering the phases associated with each episode (using the MATLAB *randperm* function). The random permutation is applied independently to each unit within each of the episodes (and hence stimuli). This last detail is significant as it abolishes phase dependencies of simultaneously recorded units. For both real and shuffled data, composite spike phase distributions of each unit are obtained by combining phase distributions across all stimuli. For statistical analyses of the phases, we used the MATLAB circ_stat toolbox (65), and specifically the following functions: *circ_otest* for uniformity of data, *circ_mean* for the circular mean. For pairwise comparisons of phase distributions, we use the *circ_cmtest* function, which is a non-parametric test that compares medians of two samples. For all our analyses, we only evaluated phase distributions (whether for individual neurons or for specific unit-stimulus combinations) if the number of spikes within the samples was at least 50.

### Analysis procedures across sessions

To statistically compare stimulus triggered averaged TMS (staTMS) across different stimuli, (as described in the text in relation to **Fig. 5** and **Fig. S7**) we combined the staTMS (across all trials for each stimulus) across all sessions. Thus, the number of data points associated with each stimulus is the number of sessions multiplied by the number of samples in each data cut. Given our LFP sampling rate of 508.63 Hz a data segment of 70 s (beginning 5 s before stimulus presentation and ending 65 after it), this yields 35605 samples. Each sample is associated with one type of stimulus, and then a non-parametric ANOVA (MATLAB *kruskalwallis* function) is used to test the null hypothesis of equal medians for all stimuli.

In several analyses (e.g., phase analyses), we associated oscillatory episodes with specific stimuli. An episode was associated with a stimulus if, and only if, it is was entirely included within a given stimulus epoch (as defined above).

To conduct pairwise comparisons of phase distributions between all pairs of neurons, we considered each pair of units. We conducted a comparison if, and only if, at least one of the two units under comparison, or their shuffled distributions, had a non-uniform phase distribution (p < 0.05 as determined using the *circ_otest*). We used this procedure because we reasoned that there is no meaning to comparing median phases of two distributions of which neither deviates from uniformity. We also considered phases of the shuffled distributions in order to make the comparison between real and shuffled distributions more balanced. If inclusion based on non-uniformity is only based on the real distribution, this presents a bias against the shuffled distributions leading to even greater differences than those observed here between the real and shuffled distributions. The same procedure was applied for comparison of phase distributions associated with pairs of stimuli for particular neurons. Specifically, we only compared pairs of phase distributions if at least one of the members (in either the real or shuffled data) significantly deviated from uniformity (p < 0.05 as determined using the *circ_otest*). We stress that our results remain entirely valid (with unequivocal differences between real and shuffled data) using other criteria for inclusion in comparisons (for example, when considering all pairs of distribution samples regardless of deviation from uniformity, or conversely, when considering only pairs in which both samples significantly deviate from uniformity).

Calculation of the probability of observing a certain number of action potentials within a window (**Fig. 8**): We used the MATLAB *binocdf* function, to calculate the upper tail of the cumulative probability to obtain x or more spikes from a population of N neurons with a certain firing rate (f). More specifically (if f is not too large), a Poisson neuron (i.e., fixed firing probability at each point in time) with a baseline rate of f Hz has a firing probability of f/1000 within a 1 ms window. Thus, the mean number of spikes emitted by n neurons within a temporal window of s ms is simply f•s•n. To quantify the effect of phase dependent firing, we compare the result when f is multiplied by various factors (1.1, 1.5, and 2).

*All the raw data, processed data, and codes will be provided by the authors upon request*.

## Acknowledgements

This work was supported by Deutsche Forschungsgemeinschaft (German Research Foundation) 378028035 to Y.B.S. and M.S and German-Israeli Foundation for Scientific Research and Development 1-1193-153.13/2012 to Y.B.S. and M.S. Israeli Science Foundation (ISF) grant (1703/16) for Y.B.S. A.K. was supported by Lady Davis postdoctoral fellowship, IL.

## Supplementary information (Table and Figures)

**Table S1:**
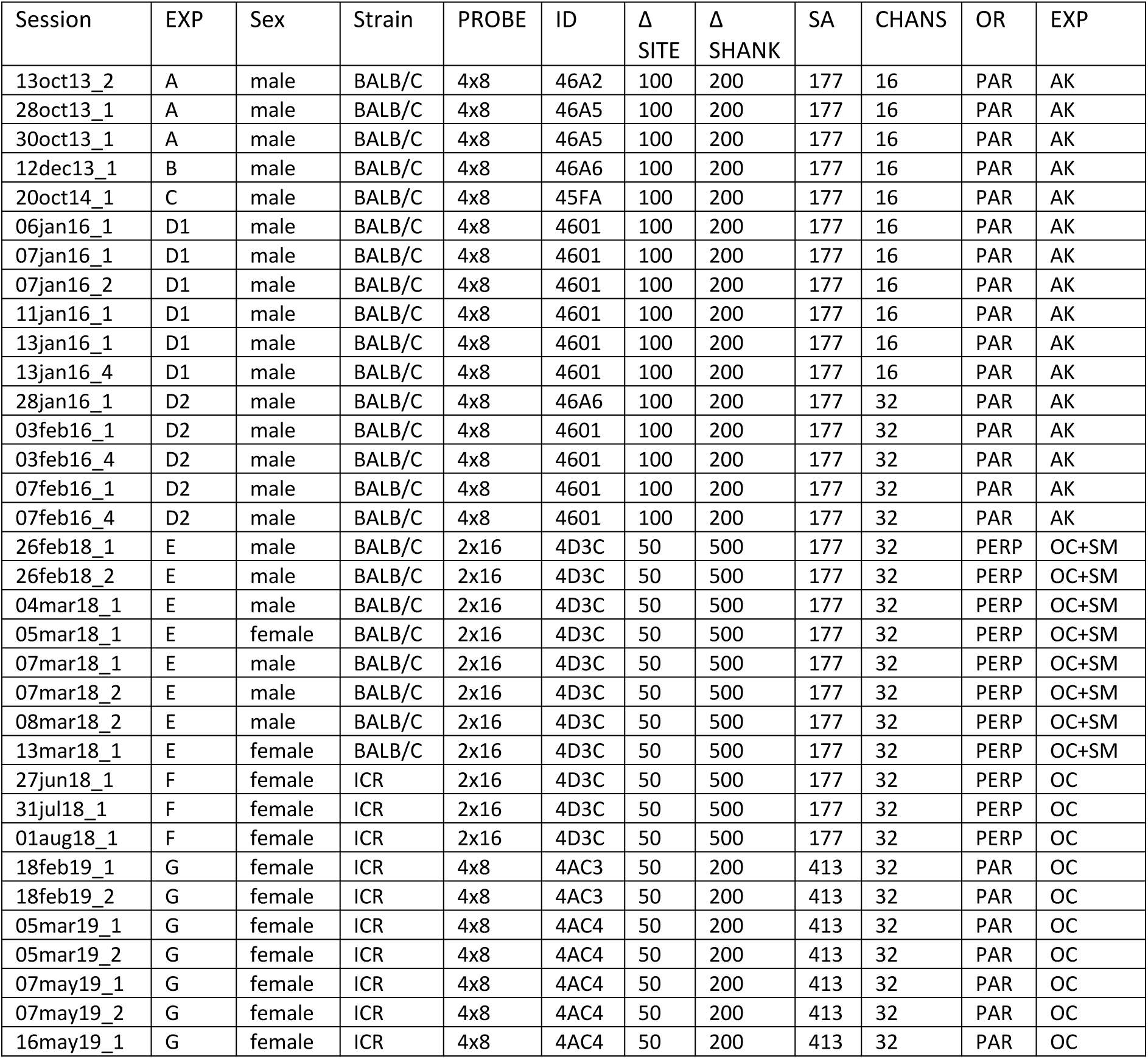
Details of the sessions used in this manuscript. The letters correspond to the experiment types indicated in the table. A. Urine, saliva, and vaginal secretions, from estrus and non-estrus female mice, from the BALB/C and C57BL/6 strains. See further details in (8). B: Urine from estrus and non-estrus female mice from the BALB/C strain at 1, 10 and 100F solution. C: Male urine, Female urine at various pH values (5–7). See (99). D1, D2: Mixtures of male, female, and predator stimuli each at 1:100, 1:30, and 1:10 dilutions in Ringer’s solution. Male and female mixtures included stimuli from the BALB/C and C57BL/6 strains. See (73). E: BALB/C female and male urine, predator urine mix, and ringer’s solution. In these experiments, we did not perform a tracheotomy, since we wanted to maintain airflow in the nasal cavity during breathing. Because of this, we could not flush the nasal cavity with solution, and thus these experiments only contain a small number of trial repeats. F: Undiluted urine mixes from castrated, dominant and naïve male mice. Each mix contains urine from the ICR, BALB/C and C57BL/6 strains. A mix of estrus and non-estrus female urine, predator urine mix, and ringer’s solution. G: Undiluted urine from naïve, dominant, and castrated male mice from the BALB/C, C57BL/6, and ICR strains (9 different stimuli), as well as urine mixes from estrus and non-estrus female mice, and a ringer’s control stimulus. Session: date and session number within date. Unique experiment ID, EXP: shorthand for type of experiment, SEX: sex of the subject mouse, STR: strain of subject mouse, PROBE: number of shanks x number of sites per shank in recording probe, ID: probe ID in the NeuroNexus catalogue, Δ SITE: vertical distance between sites on the same shank (µm). Δ SHANK: distance between shanks on the probe (µm). SA: site area (µm^2^), CHANS: number of active LFP channels, OR: probe orientation. PAR: parallel to the AOB-ECL, PERP: perpendicular to the AOB-ECL. EXP: initials of lead experimenter.

| Session | EXP | Sex | Strain | PROBE | ID | $\Delta$ SITE | $\Delta$ SHANK | SA | CHANS | OR | EXP |
| --- | --- | --- | --- | --- | --- | --- | --- | --- | --- | --- | --- |
| 13oct13_2 | A | male | BALB/C | 4x8 | 46A2 | 100 | 200 | 177 | 16 | PAR | AK |
| 28oct13_1 | A | male | BALB/C | 4x8 | 46A5 | 100 | 200 | 177 | 16 | PAR | AK |
| 30oct13_1 | A | male | BALB/C | 4x8 | 46A5 | 100 | 200 | 177 | 16 | PAR | AK |
| 12dec13_1 | B | male | BALB/C | 4x8 | 46A6 | 100 | 200 | 177 | 16 | PAR | AK |
| 20oct14_1 | C | male | BALB/C | 4x8 | 45FA | 100 | 200 | 177 | 16 | PAR | AK |
| 06jan16_1 | D1 | male | BALB/C | 4x8 | 4601 | 100 | 200 | 177 | 16 | PAR | AK |
| 07jan16_1 | D1 | male | BALB/C | 4x8 | 4601 | 100 | 200 | 177 | 16 | PAR | AK |
| 07jan16_2 | D1 | male | BALB/C | 4x8 | 4601 | 100 | 200 | 177 | 16 | PAR | AK |
| 11jan16_1 | D1 | male | BALB/C | 4x8 | 4601 | 100 | 200 | 177 | 16 | PAR | AK |
| 13jan16_1 | D1 | male | BALB/C | 4x8 | 4601 | 100 | 200 | 177 | 16 | PAR | AK |
| 13jan16_4 | D1 | male | BALB/C | 4x8 | 4601 | 100 | 200 | 177 | 16 | PAR | AK |
| 28jan16_1 | D2 | male | BALB/C | 4x8 | 46A6 | 100 | 200 | 177 | 32 | PAR | AK |
| 03feb16_1 | D2 | male | BALB/C | 4x8 | 4601 | 100 | 200 | 177 | 32 | PAR | AK |
| 03feb16_4 | D2 | male | BALB/C | 4x8 | 4601 | 100 | 200 | 177 | 32 | PAR | AK |
| 07feb16_1 | D2 | male | BALB/C | 4x8 | 4601 | 100 | 200 | 177 | 32 | PAR | AK |
| 07feb16_4 | D2 | male | BALB/C | 4x8 | 4601 | 100 | 200 | 177 | 32 | PAR | AK |
| 26feb18_1 | E | male | BALB/C | 2x16 | 4D3C | 50 | 500 | 177 | 32 | PERP | OC+SM |
| 26feb18_2 | E | male | BALB/C | 2x16 | 4D3C | 50 | 500 | 177 | 32 | PERP | OC+SM |
| 04mar18_1 | E | male | BALB/C | 2x16 | 4D3C | 50 | 500 | 177 | 32 | PERP | OC+SM |
| 05mar18_1 | E | female | BALB/C | 2x16 | 4D3C | 50 | 500 | 177 | 32 | PERP | OC+SM |
| 07mar18_1 | E | male | BALB/C | 2x16 | 4D3C | 50 | 500 | 177 | 32 | PERP | OC+SM |
| 07mar18_2 | E | male | BALB/C | 2x16 | 4D3C | 50 | 500 | 177 | 32 | PERP | OC+SM |
| 08mar18_2 | E | male | BALB/C | 2x16 | 4D3C | 50 | 500 | 177 | 32 | PERP | OC+SM |
| 13mar18_1 | E | female | BALB/C | 2x16 | 4D3C | 50 | 500 | 177 | 32 | PERP | OC+SM |
| 27jun18_1 | F | female | ICR | 2x16 | 4D3C | 50 | 500 | 177 | 32 | PERP | OC |
| 31jul18_1 | F | female | ICR | 2x16 | 4D3C | 50 | 500 | 177 | 32 | PERP | OC |
| 01aug18_1 | F | female | ICR | 2x16 | 4D3C | 50 | 500 | 177 | 32 | PERP | OC |
| 18feb19_1 | G | female | ICR | 4x8 | 4AC3 | 50 | 200 | 413 | 32 | PAR | OC |
| 18feb19_2 | G | female | ICR | 4x8 | 4AC3 | 50 | 200 | 413 | 32 | PAR | OC |
| 05mar19_1 | G | female | ICR | 4x8 | 4AC4 | 50 | 200 | 413 | 32 | PAR | OC |
| 05mar19_2 | G | female | ICR | 4x8 | 4AC4 | 50 | 200 | 413 | 32 | PAR | OC |
| 07may19_1 | G | female | ICR | 4x8 | 4AC4 | 50 | 200 | 413 | 32 | PAR | OC |
| 07may19_2 | G | female | ICR | 4x8 | 4AC4 | 50 | 200 | 413 | 32 | PAR | OC |
| 16may19_1 | G | female | ICR | 4x8 | 4AC4 | 50 | 200 | 413 | 32 | PAR | OC |

**Fig. S1.**
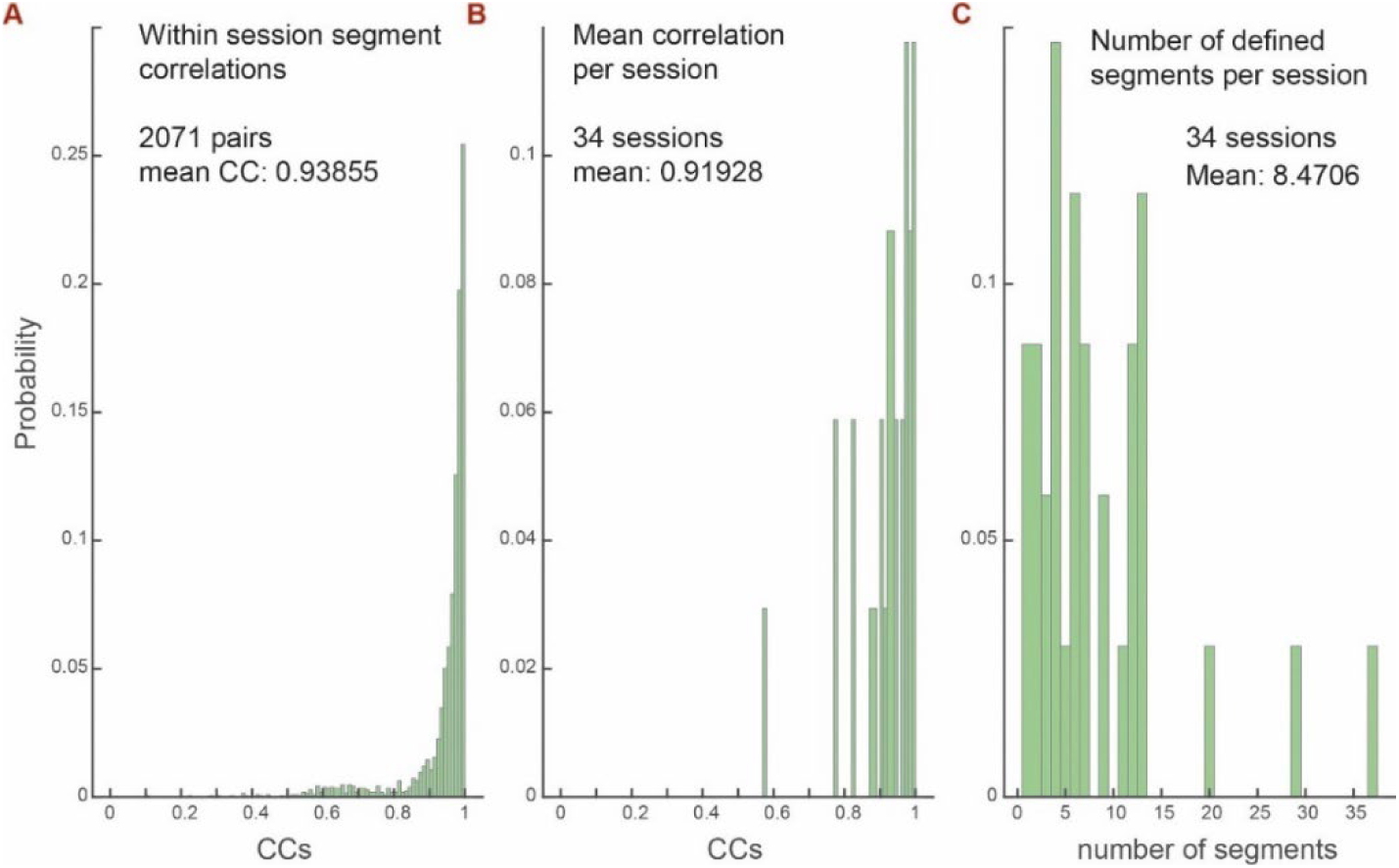
Features of manually defined events (“seed” events). **A.** Distribution of cross correlation coefficients among all manually defined events within a session, pooled over all sessions. **B.** Distribution of the mean cross correlations among events, for each session. **C**. Distribution of the number of manually defined events per session.

**Fig. S2.**
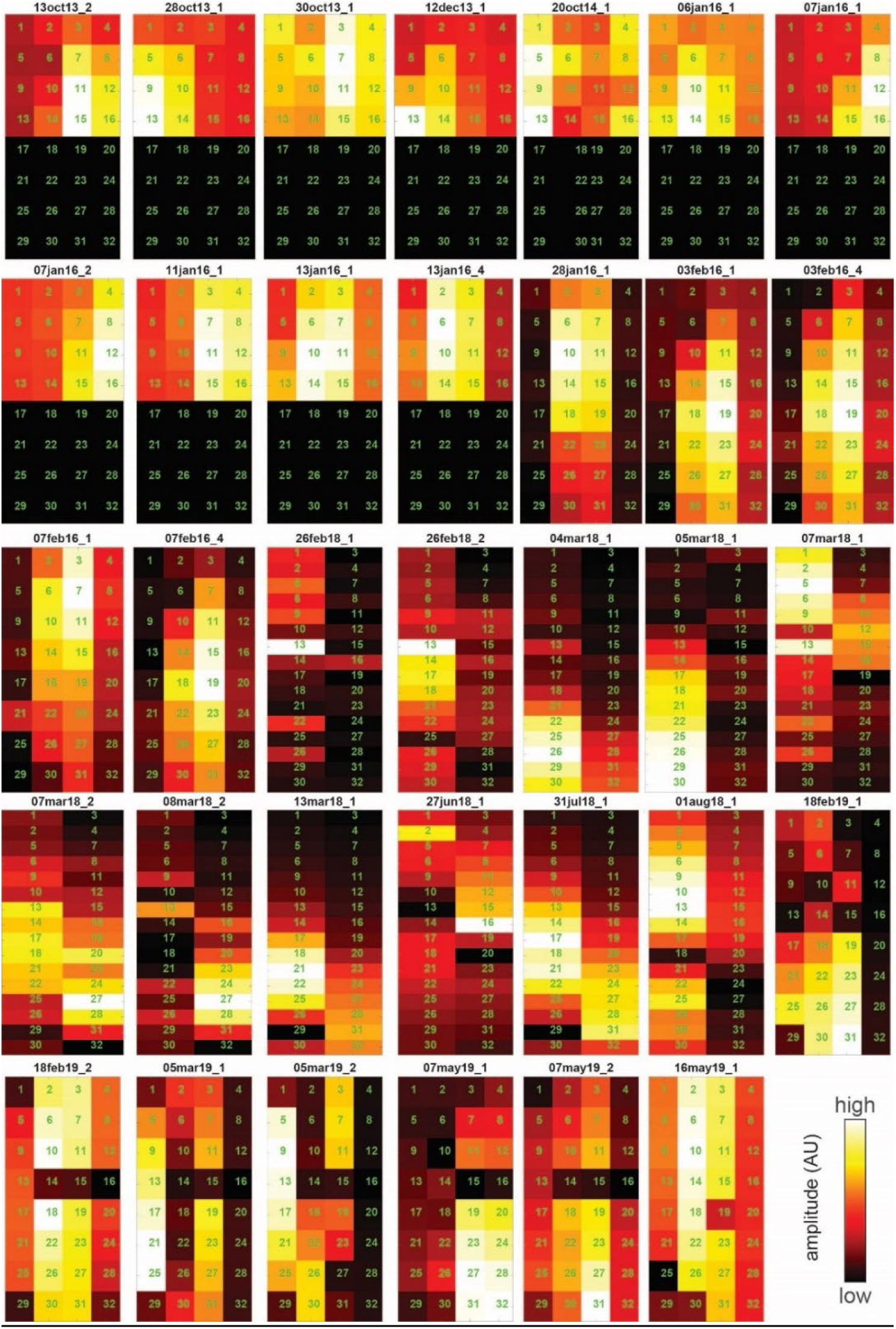
Templates for each of the sessions in the database. Note that the first 11 recordings only include 16 channels. Note also that some recordings contain non-functional recording sites. (i.e., 14-16) in some of the recordings in the lowest row. These broken channels were excluded for template calculation.

**Fig. S3.**
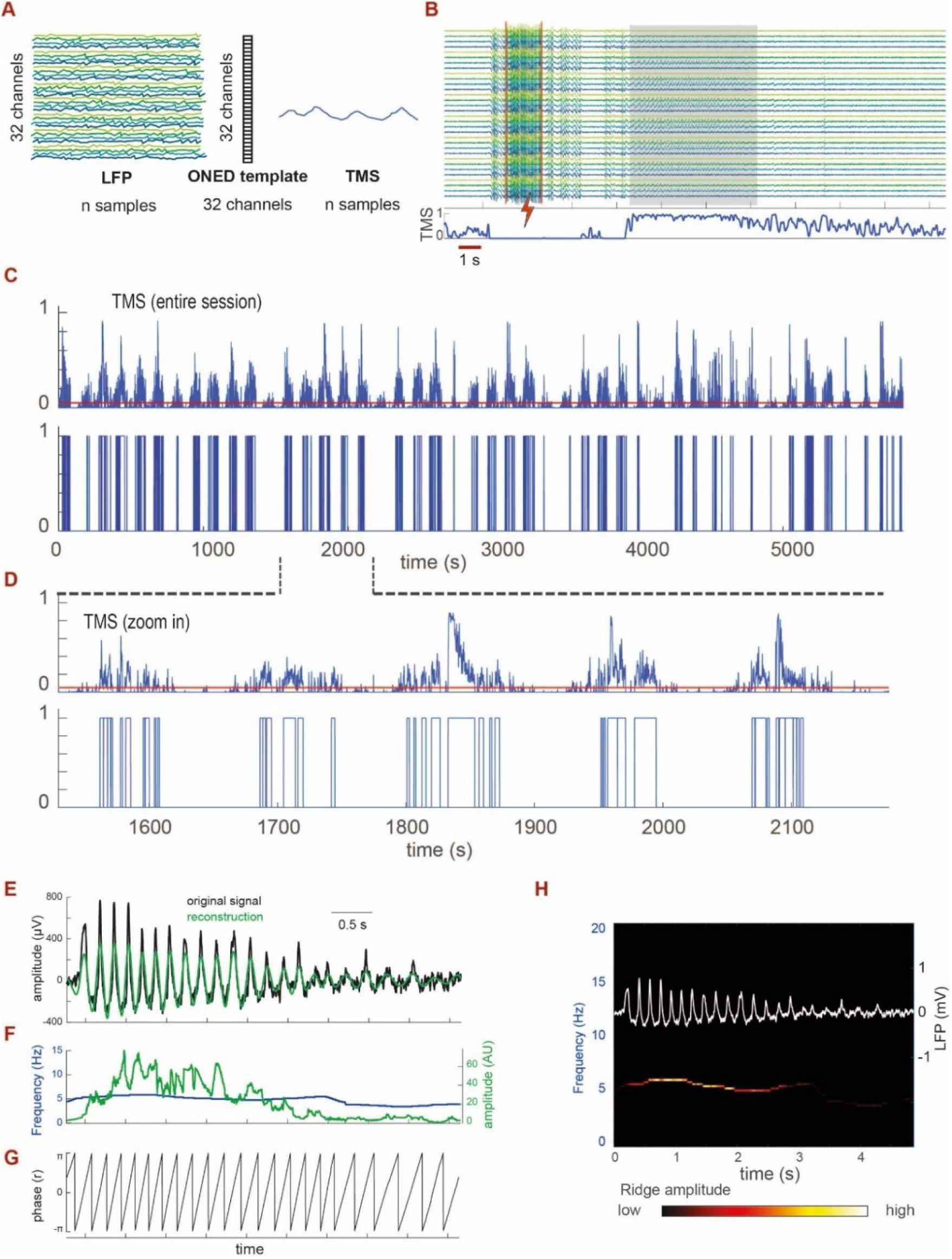
Detection of oscillatory segments and the template match approach. **A.** Schematic view of the template match signal (TMS) derivation. The signal is derived by correlating the 32 channel LFP envelope at each time point with the oscillatory network energy distribution template (ONED, as shown in Fig. 2F, G). The result is smoothed and raised to the 10th power to increase contrast (see Methods). **B.** Example of the TMS (blue trace at bottom), shown for the same segment shown in Fig. 2B. Note that the TMS is highest during the oscillatory event (shaded rectangle), and low during the period associated with VNO stimulation and with other “noisy” segments. Notably, although the TMS decreases after the oscillatory event, it remains positive even after the oscillations have subsided, reflecting the relative distribution of LFP envelopes. **C**. TMS (top) and binary above-threshold segments (bottom) during an entire session. The threshold is shown in red. **D**. Expanded scale of a section from panel C. **E** An oscillatory event in one channel showing the original signal (black) and its reconstruction from a single ridge (green). **F.** Time evolving frequency (blue) and amplitude (green) of the first ridge. **G.** Instantaneous phase of the first ridge. **H.** The original signal and the first ridge spectrogram. In this representation the ridge amplitude (green trace in panel F) is indicated by heatmap color.

**Fig. S4.**
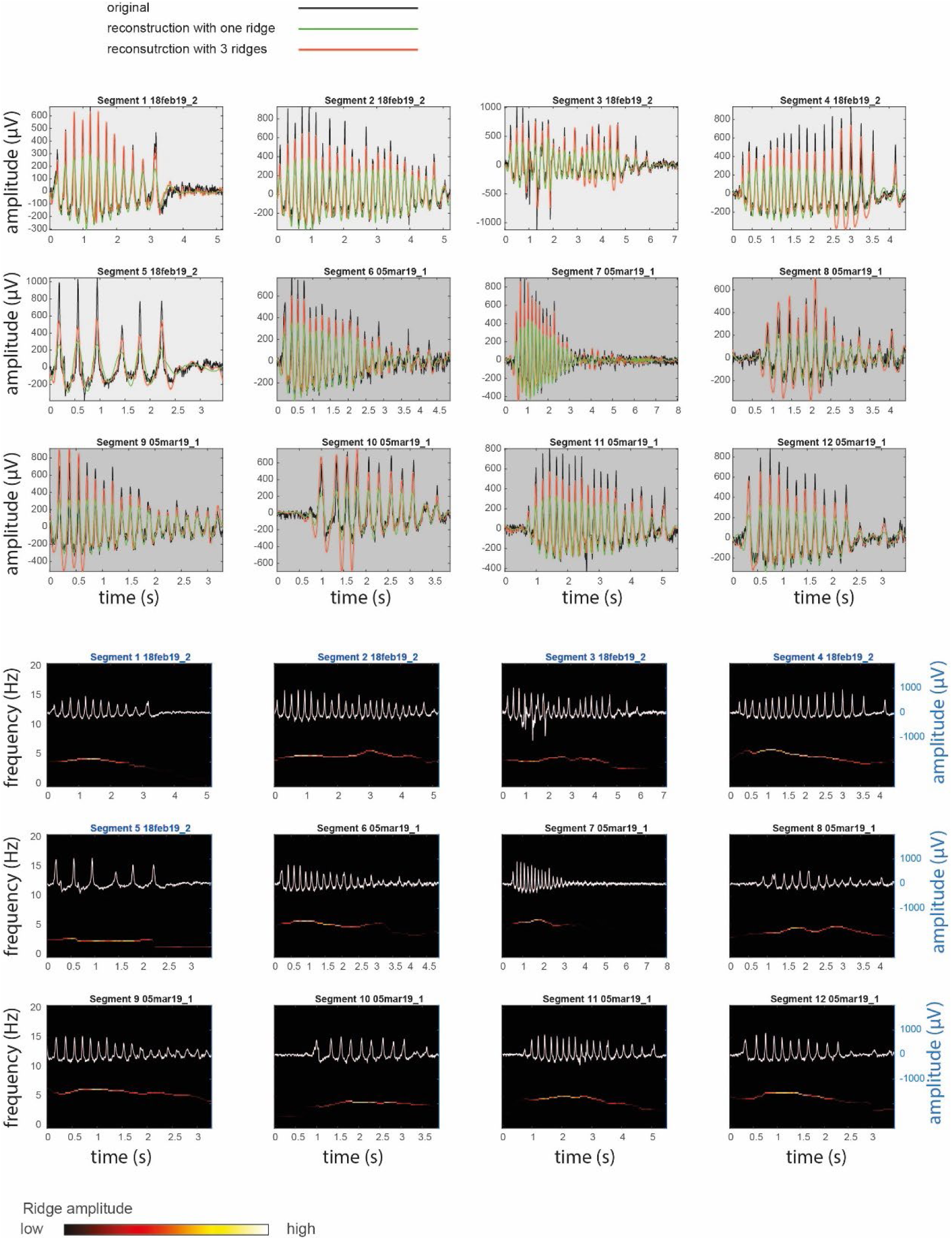
Complete episodes and their reconstructions. Each episode is shown twice. The top representation shows the original episode (black), and its reconstruction using one (green), or all 3 ridges (red). The bottom representation shows the original episode (μV scale on the right) and the first ridge time frequency decomposition as a heat map. Consecutive episodes with the same background shade were recorded in the same session (i.e., the top row and the first example on the second row are from the same session). This is one of 8 figures which together show all complete episodes. The session from which the episodes is derived in shown above each panel (shown is one panel of eight).

**Fig. S5.**
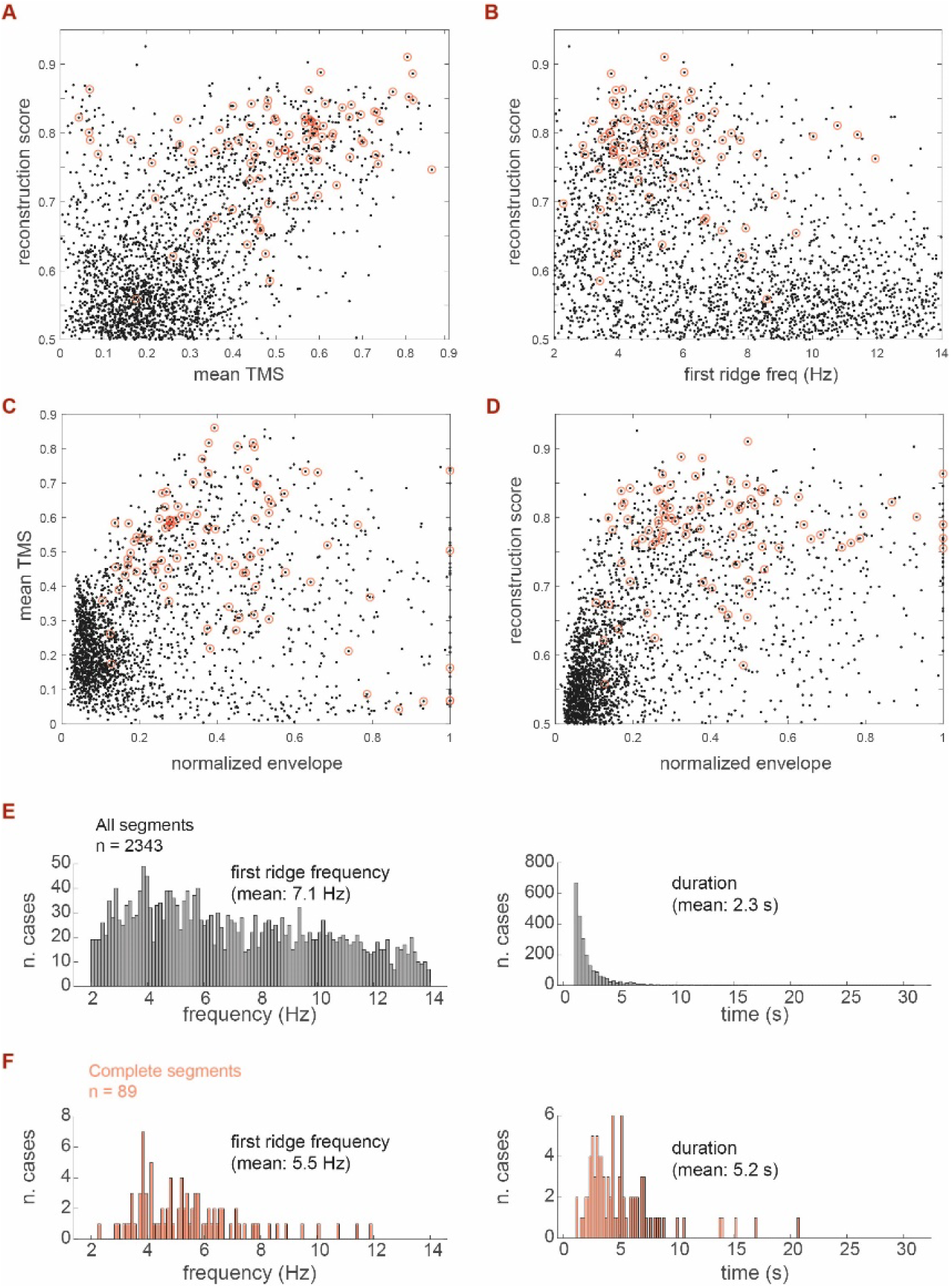
General episode features. **A**. Reconstruction score (correlation coefficient between original signal and reconstruction with the first ridge) vs. mean TMS (across the entire episode). **B.** Reconstruction score vs. the mean first ridge frequency. **C**. Mean TMS vs. normalized envelope amplitude (envelope within each session was normalized by the envelope of the episode with the largest envelope). **D**. Reconstruction score as a function of normalized envelope. In all panels, black dots represent all episodes, and black dots with red circles correspond to *complete* episodes. Note the correlation between mean template match and an above threshold reconstruction score, and the concentration of first ridge frequencies in the ∼5Hz for the complete episodes. **E.** Distribution of mean (first ridge) frequency and duration for all episodes (excluding false positives). **F.** Distribution of frequencies and durations for complete episodes. Same plots as in Fig. 3B, C, included here for comparison.

**Fig. S6.**
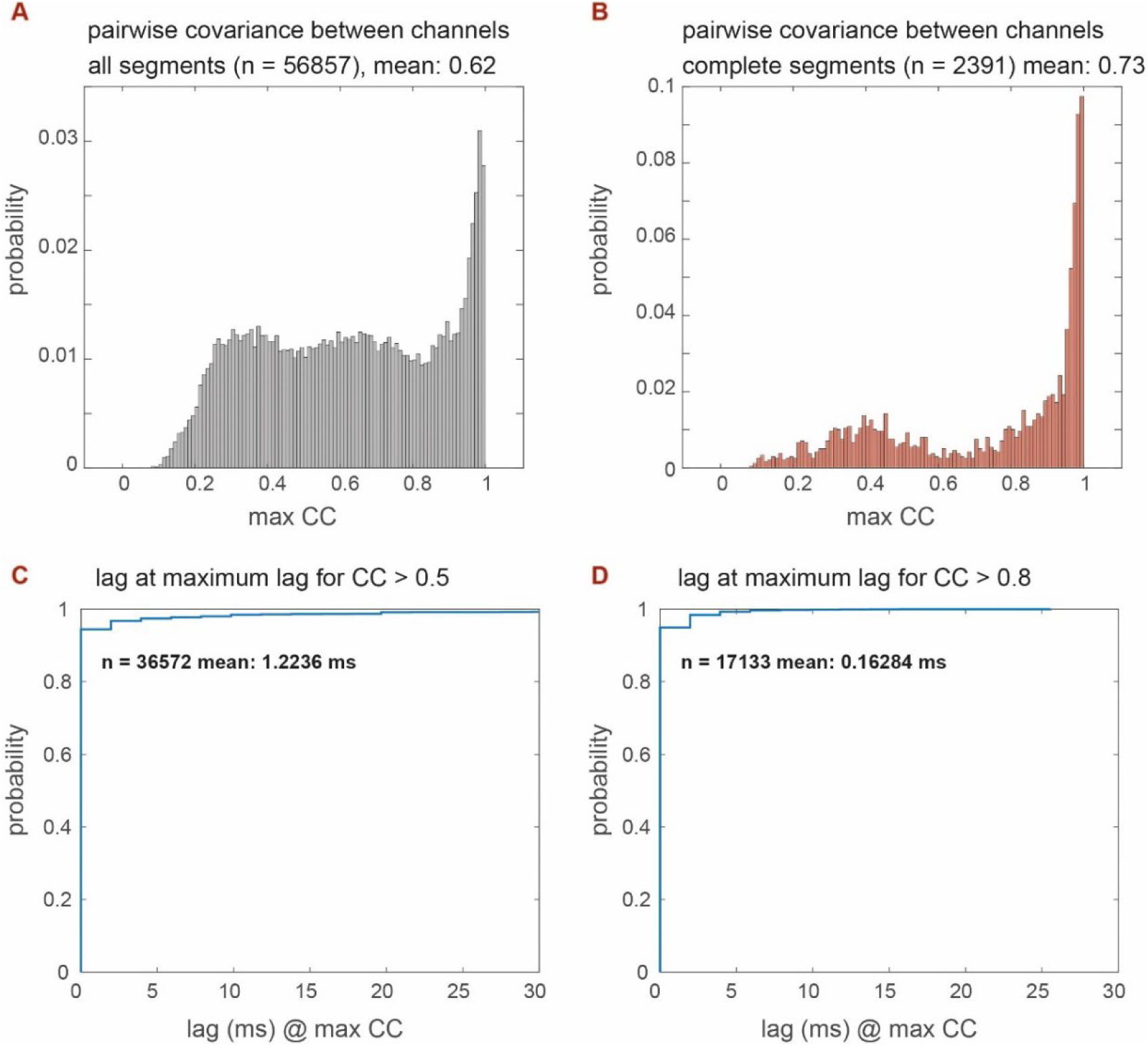
Analysis of temporal lags across channels within an episode. For each episode, we used the cross-covariance (CC) function (*xcov* function in MATLAB) to find the best match (highest covariance) between the channel with the maximum energy (for that episode), and all other channels. **A.** Distribution of all CC obtained with this procedure. **B.** Distributions of CCs for *complete* episodes. **C.** Cumulative distribution of lags (in absolute values) that yield the highest CC. To exclude non-oscillating channels from this analysis, we only considered channels that had a CC of at least 0.5 (a low threshold considering the distribution shown in A). The rationale for setting a threshold on the CC is that non-oscillating channels will not yield high correlations with the highest-amplitude, reference oscillating channel **D.** Cumulative lag distributions when considering channels with CCs of at least 0.8. Mean values are indicated in panels C and D and are very small relative to a theta cycle. This analysis revealed that percentage of channels with lags smaller than 10 ms (in absolute value) are 85% across all channels in all episodes (no CC threshold), 98% for all channels with a CC > 0.5 (as in panel C), and 99.9% when considering channels with CC > 0.8 (panel D). A period of 10 ms represents 1/20 of a 5 Hz theta cycle and thus assigning the oscillation phase based on one channel is expected to induce a negligible jitter in our phase estimates.

**Fig. S7.**
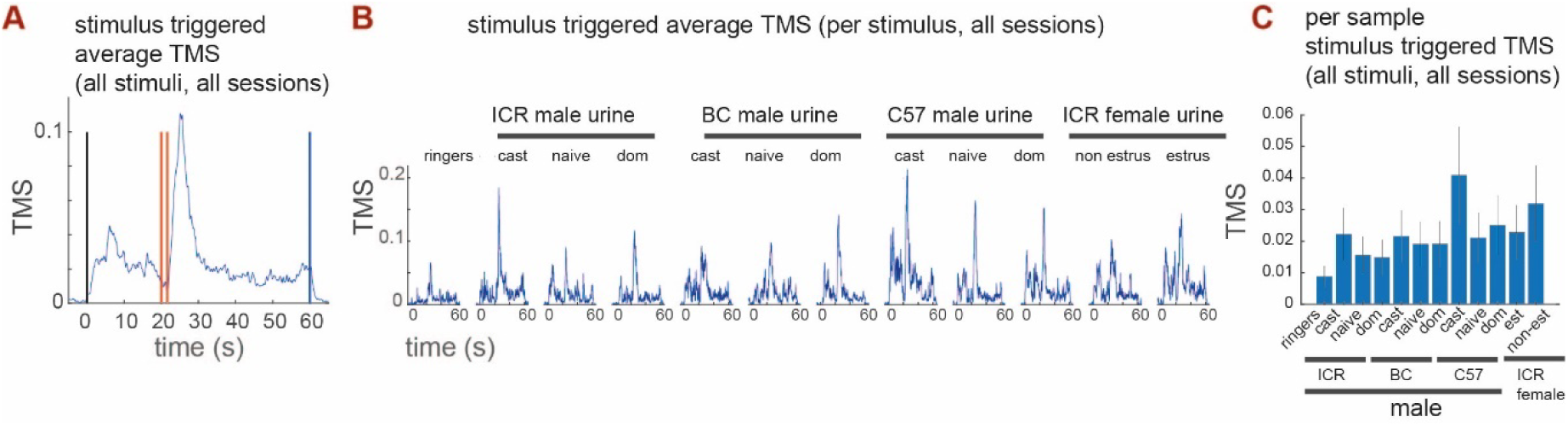
The mean TMS, around all stimulus presentation times, in one type of experiment. Like Fig. 5B**-D**, but for a different stimulus set (exp G, **Table S1**)**. A.** Stimulus triggered average TMS (staTMS), around all stimulus presentation times, in one type of experiment. **B.** Average TMS for each stimulus separately. **C.** The mean template signal (directly proportional to areas in B) and the standard error across all presentations of each of the stimuli. See Table S1 for details on the stimuli used. Note that different stimuli elicit different TMS profiles, and that the control stimulus (ringers) induces the weakest response.

**Fig. S8.**
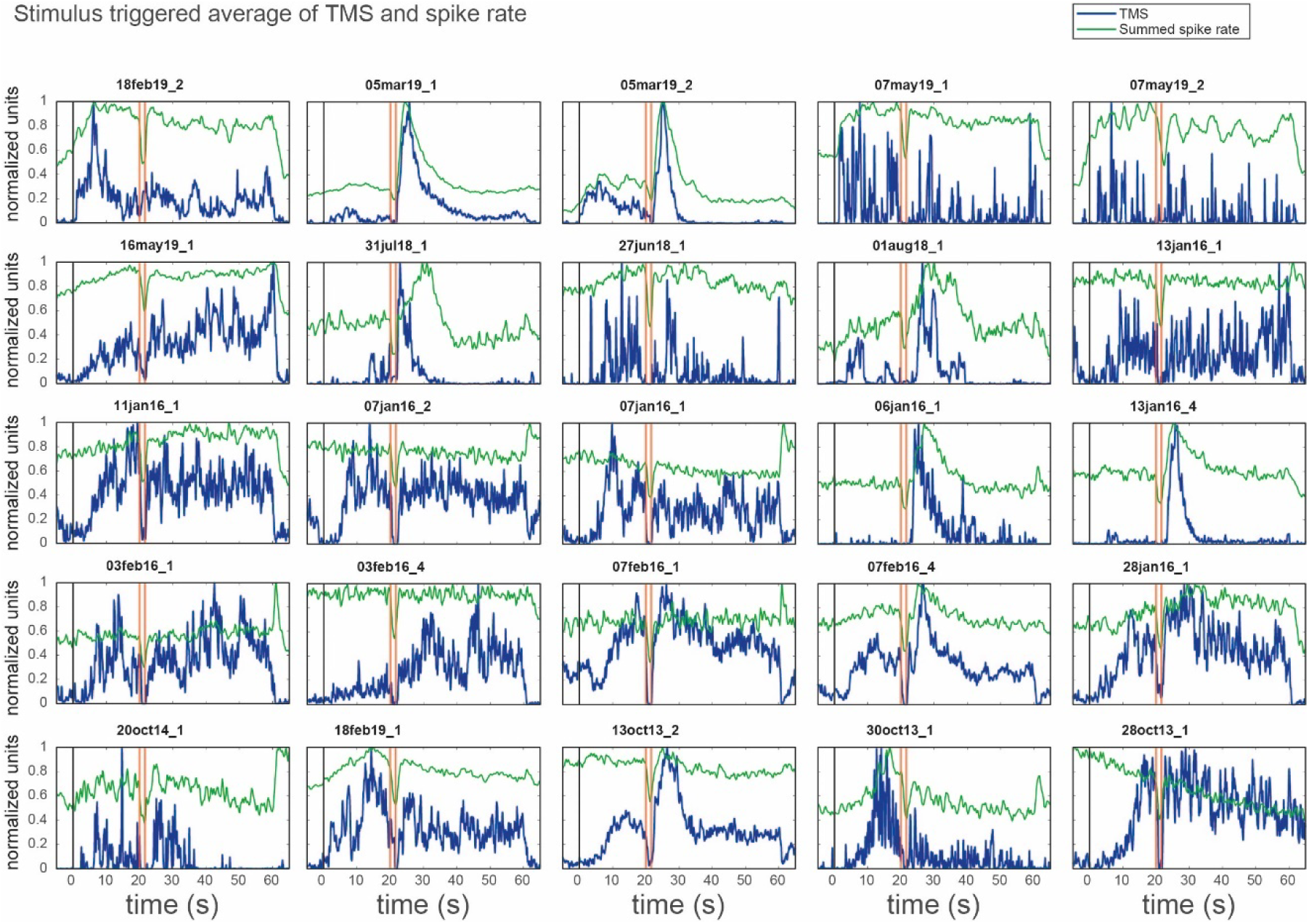
Relationship between the TMS and summed single unit activity, across stimulus presentations. Same conventions as **Fig. 6C**, but with more sessions included. Stimulus triggered averages of both values were calculated across all stimuli. Green and blue traces correspond to the spike signal, and the TMS, respectively. Note that the TMS generally follows the stimulation time course more closely and is more localized in time than the spike signal.

**Fig. S9.**
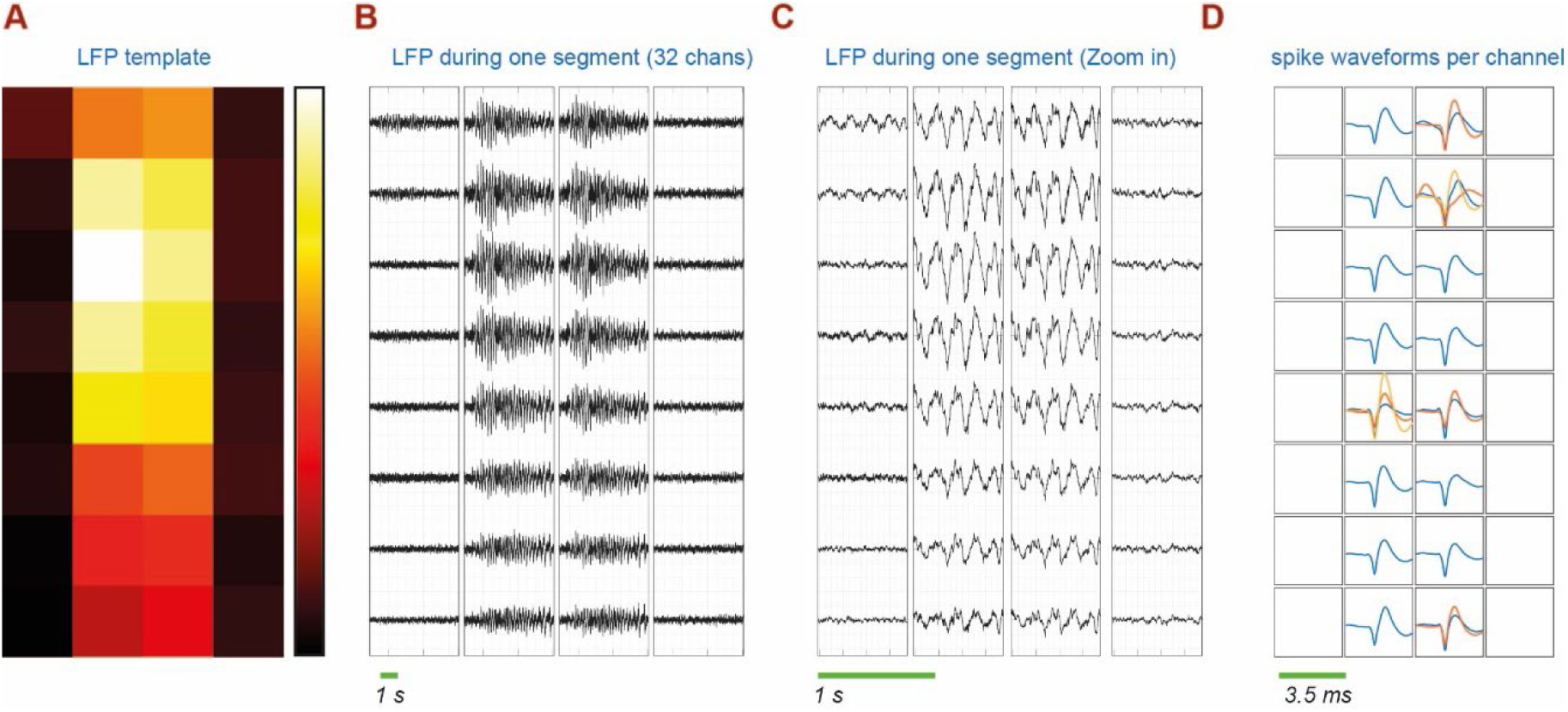
Evidence for localization of oscillations within the AOB ECL. Like Fig. 7C**-F**, but for a different session. **A.** Spatial spread of LFP energies in one of the sessions. **B.** LFP oscillations for one oscillating episode, across all channels, plotted according to spatial position. **C.** Magnified temporal view of B, highlighting the phase similarity across all channels. **D**. Spike waveforms in the sites shown in A. Each square shows the spike waveforms recorded in that channel. Both single unit and multi-unit activity waveforms are shown.

